# Evidence for multifactorial processes underlying phenotypic variation in bat visual opsins

**DOI:** 10.1101/300301

**Authors:** Alexa Sadier, Kalina T. J. Davies, Laurel R. Yohe, Kun Yun, Paul Donat, Brandon P. Hedrick, Elizabeth R. Dumont, Liliana M. Dávalos, Stephen J. Rossiter, Karen E. Sears

**Author notes:** Co-first authors (Alexa Sadier, Kalina T. J. Davies). Co-corresponding authors: Karen E. Sears, Ecology and Evolutionary Biology Department, UCLA, Los Angeles, CA, Stephen Rossiter, School of Biological and Chemical Sciences, Queen Mary University of London, London, United Kingdom, Liliana M. Dávalos, Department of Ecology and Evolution, Stony Brook University, Stony Brook, New York, USA.

## Abstract

Studies of opsin genes offer insights into the evolutionary history and molecular basis of vertebrate color vision, but most assume intact open reading frames equate to functional phenotypes. Despite known variation in opsin repertoires and associated visual phenotypes, the genetic basis of such patterns has not been examined at each step of the central dogma. By comparing sequences, gene expression, and protein localization across a hyperdiverse group of mammals, noctilionoid bats, we find evidence that independent losses of S-opsin arose through disruptions at different stages of protein synthesis, while maintenance relates to frugivory. Discordance between DNA, RNA, and protein reveals that the loss of short-wave sensitivity in some lineages resulted from transcriptional and post-transcriptional changes in addition to degradation of open reading frames. These mismatches imply that visual phenotypes cannot reliably be predicted from genotypes alone, and connect ecology to multiple mechanisms behind the loss of color in vertebrates.

## Introduction

Studies of opsin gene sequences, which encode the photoreceptor proteins within rod and cone cells, have been crucial to understanding the molecular basis and evolutionary history of visual perception in animals, and provide some of the most compelling examples of how substitutions in the genetic code can bring about adaptive phenotypic change (Lucas et al., 2003; Porter et al., 2012; Yokoyama et al., 2008). In particular, comparisons of amino acids thought to underpin spectral tuning help infer the visual abilities and sensory ecologies of extant taxa and, through ancestral sequence reconstructions, extinct hypothetical forms (Hunt et al., 2009; Zhao, Rossiter, et al., 2009). Other studies have revealed allelic variation in opsin genes within populations (Jacobs et al., 2017), and the contribution of duplications (Cortesi et al., 2015; Feuda et al., 2016) and adaptive gene losses (Davies et al., 2009) to photoreceptor repertoires.

Inferring phenotype solely from gene sequence, however, rests on the assumption that genetic information is invariably transcribed from DNA to RNA, and translated from RNA to protein in the presence of an intact open reading frame. However, while a growing number of studies have examined the expression of opsin genes (Cheng & Flamarique, 2004) and/or their protein products (Peichl et al., 2017; Wikler & Rakic, 1990), no comparative studies of opsins have probed each step of this central dogma. Thus, the extent to which visual phenotypes are expressed or masked due to the modulation of protein production is currently unknown. This represents a major gap in our understanding of visual evolution, as mounting evidence from a range of systems is beginning to reveal that complex post-transcriptional and post-translational routes shape phenotypic variation and complicate genotype-to-phenotype mapping (Blount et al., 2012; Csardi et al., 2015; Schwanhausser et al., 2011).

The potential for selection to act on phenotypes via multiple molecular mechanisms may be particularly important in rapid functional trait diversification, as is often the case in visual systems. In sticklebacks, for example, the repeated colonization of lakes with different photopic environments has driven shifts in spectral sensitivity via recurrent selective sweeps in short-wave opsin genes, and changes in opsin expression (Marques et al., 2017; Rennison et al., 2016). Similarly, rapid shifts in the visual ecology of cichlid fishes have involved a combination of coding sequence evolution and changes in expression (O’Quin et al., 2010; Spady et al., 2005). In contrast, much less is known about the mechanisms underpinning rapid visual adaptations in mammals and reptiles, for which relevant studies have tended to focus on ancient transitions to nocturnal, aquatic or subterranean niches (Emerling, 2017; Emerling et al., 2017; Jacobs et al., 1993).

Bats, like most mammals, possess functional open reading frames of *rhodopsin* (*RHO*) (Zhao, Ru, et al., 2009), *opsin 1 long-wave sensitive* (*OPN1LW*), and, in most cases, *opsin 1 short-wave sensitive* (*OPN1SW*) genes (Zhao, Rossiter, et al., 2009). A few Old World bat lineages are known to have lost their short-wave opsins, but they are all confined to the suborder Yinpterochiroptera (Zhao, Rossiter, et al., 2009). Immunohistochemical screens of rods and cones in New and Old World bats have thus far supported these trends, albeit based on very few taxa (Butz et al., 2015; Feller et al., 2009; Kim et al., 2008; Müller et al., 2009; Müller et al., 2007). Inferences from amino acid sequence analyses and action spectra suggest that bat short-wave opsins are sensitive to UV, and their retention is possibly related to the demands of visual processing in mesopic, or low-light, conditions (Zhao, Rossiter, et al., 2009), and/or plant visiting (Butz et al., 2015; Feller et al., 2009; Kim et al., 2008; Müller et al., 2009; Müller et al., 2007). However, limited taxonomic sampling precludes clear conclusions.

Bats of the superfamily Noctilionoidea (∼200 species of New World Leaf-nosed bats and allies within the suborder Yangochiroptera) underwent unparalleled ecological diversification starting about 40 million years ago (Dumont et al., 2012; Rojas et al., 2012; Rossoni et al., 2017). Noctilionoids exhibit marked morphological (Dumont et al., 2012; Monteiro & Nogueira, 2010) and sensory (Hayden et al., 2014; Yohe et al., 2017) adaptations that complement an impressive diversity of dietary niches, making them an outstanding group of mammals in which to examine the genetic and developmental basis of visual adaptations. In noctilionoid bats, switches in feeding ecology from generalized insectivory to blood-, insect-, vertebrate-, nectar- or fruit-based diets have occurred multiple times among closely related species. To determine whether dietary shifts in noctilionoid bats are associated with molecular adaptations in their visual systems, we applied analyses of sequence evolution, gene expression, and immunohistochemistry across the taxonomic and ecological breadth of the clade and outgroup taxa. By assessing the correspondence between the presence or absence of a functional open reading frame, its transcript, and its protein product for the two cone opsins (OPN1SW and OPN1LW), we identify specific and diverse molecular mechanisms by which selection has acted. Thus, by accounting for all the steps of genotype-to-phenotype correspondence, we characterize the evolutionary history of color vision in a hyperdiverse group and the contribution of ecological selective demands to shaping this evolutionary history.

## Results

We used complementary approaches to determine the distribution of color vision in noctilionoid bats. First, we performed RNA-Seq to assess the presence or absence of transcripts for *OPN1SW*, *OPN1LW*, and *RHO*, and estimated the mode and the strength of selection in coding sequences. Second, for the same taxa, we used immunohistochemistry to characterize and quantify S- and L-opsin proteins in the retinas of adult bats. Finally, we modeled the presence or absence of S-opsin cones, as indicated by protein presence, as a function of dietary ecology.

### Variation in opsin transcripts across taxa

We generated eye transcriptomes for 38 species and detected the presence of *OPN1SW* transcripts in 32 of these (see Figure 1 and Table S1). In comparison, we recovered the complete *RHO* and *OPN1LW* transcript from all taxa (*n* = 38; Figure 1). The absence of the *OPN1SW* transcript sequence was phylogenetically widespread (Figure 1). In most species examined, the ATG start codon appears to occur three codons downstream relative to that of the human orthologue. Close inspection of transcript sequences revealed an in-frame three base pair deletion (Y190del) in three of four *Pteronotus parnellii* individuals sequenced, as well as a premature stop codon in *Carollia brevicauda* that is indicative of a transcribed pseudogene. Finally, we detected multiple introns in the assembled transcripts of the sister taxa *Erophylla bombifrons* and *Phyllonycteris poeyi*, suggesting transcriptional readthrough.

**Figure 1.**
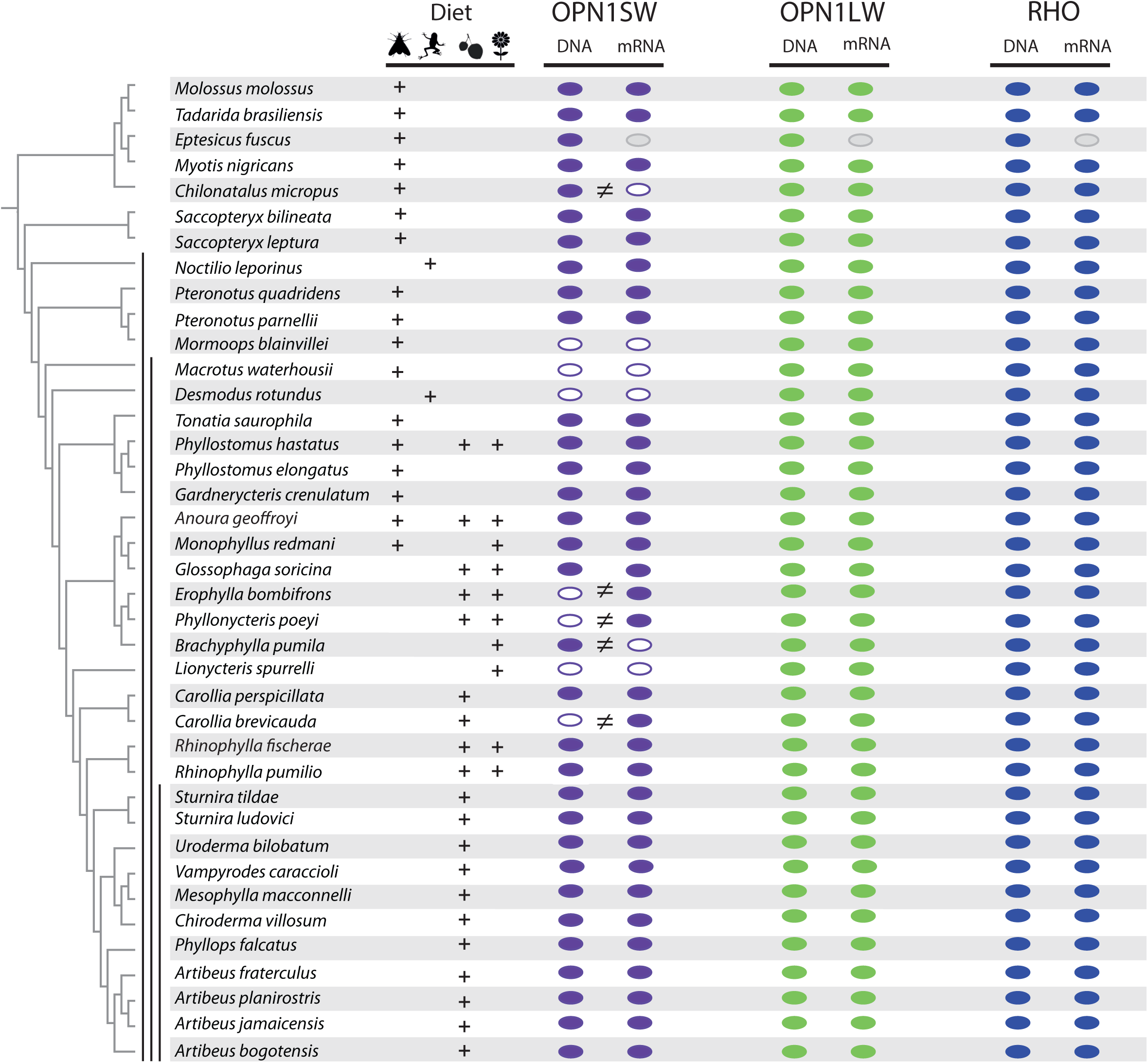
Distribution of an intact open reading frame (ORF) and mRNA transcript for the OPN1SW, OPN1LW and RHO photopigments in ecologically diverse noctilionoid bats. The composition of species diet follows Rojas et al. (2018), dietary types are indicated with the following symbols: invertebrates – moth, vertebrates – frog, fruit – fruit and nectar/pollen – flower. The species phylogeny follows (Rojas et al., 2016; Shi & Rabosky, 2015). Vertical black bars, from left to right, indicate: (1) Noctilionoidea, (2) Phyllostomidae, (3) and Stenodermatinae, respectively. RNA-Seq data was generated to both infer the presence of an intact ORF (in combination with genomic and PCR sequence data) and determine the presence of an expressed mRNA transcript. The presence of an intact ORF and mRNA transcript for *RHO* was verified across all transcriptomes. The presence of an intact ORF, mRNA and protein are indicated by a filled color marker (*OPN1SW*– purple, *OPN1LW*– green and *RHO* – blue), and its absence by a white marker. Missing data (i.e. species for which we were unable to obtain tissue) are indicated with a grey marker. Mismatches between intact ORFs and transcripts are indicated by an inequality symbol.

### Variation in opsin repertoires

We used immunohistochemistry to assess the presence or absence of OPN1SW and OPN1LW proteins in whole, flat-mounted retinas of adult bats (*n_eyes_* = 218, *n_individuals_* = 187, *n_species_* = 56). OPN1SW was detected in 32 species, but was absent in 18 species. In contrast, we confirmed the presence of OPN1LW protein in every species examined (Figure 2 and Figure supplement 1). Samples derived from museum specimens taken from six additional species had low signal-to-background ratios in the OPN1SW protein labeling, generating inconclusive results. We then analyzed the patterns of OPN1SW and OPN1LW protein expression among cells. Consistent with cone-specific roles, we found that almost all cones expressed either OPN1SW or OPN1LW protein exclusively, with no strong evidence of co-localization of both proteins (Figure 3 and Figure supplement 1).

**Figure 2.**
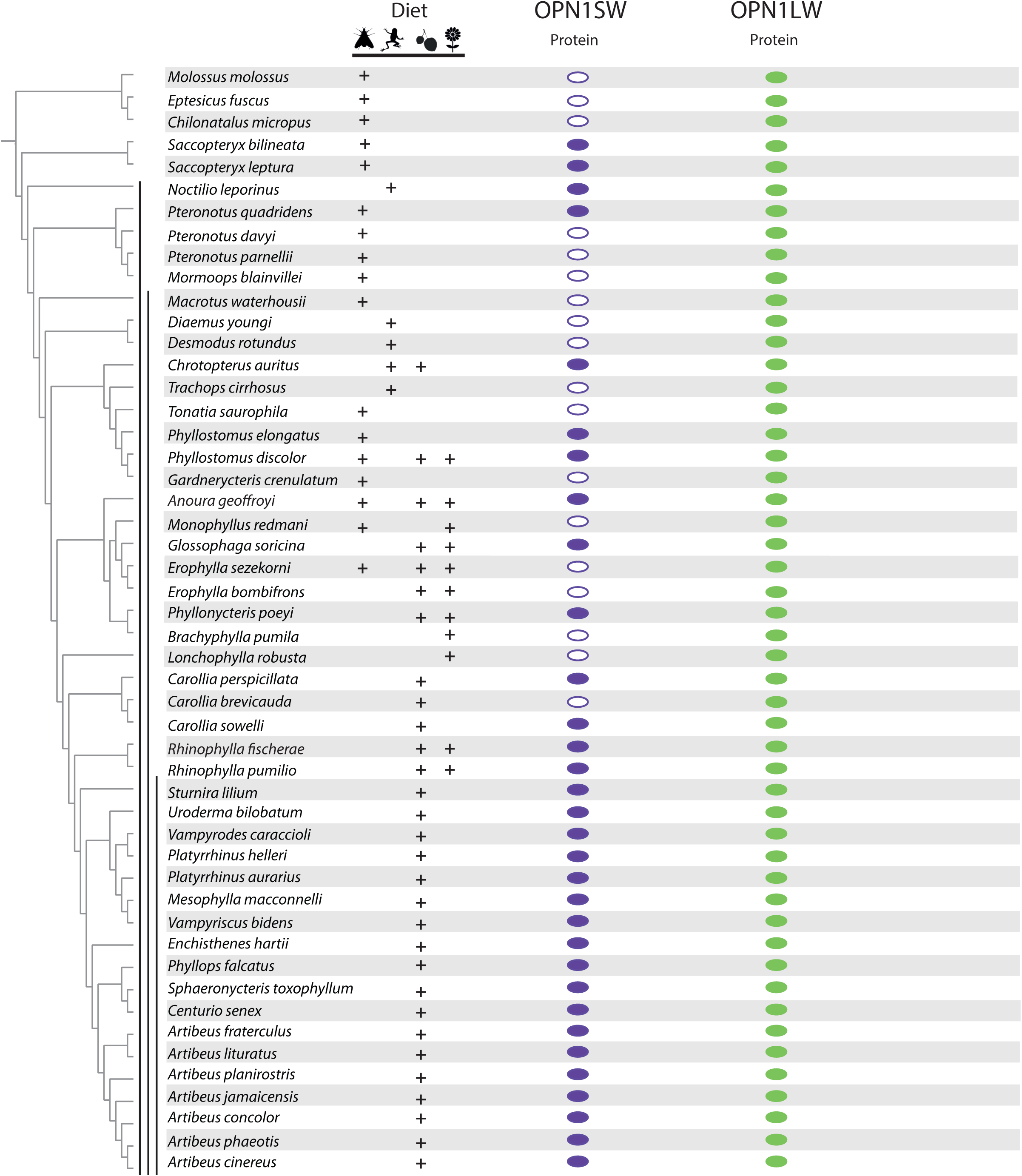
Distribution of protein for the OPN1SW and OPN1LW photopigments in ecologically diverse noctilionoid bats. The composition of species diet follows Rojas et al. (2018), dietary types are indicated with the following symbols: invertebrates – moth, vertebrates – frog, fruit – fruit and nectar/pollen – flower. The species phylogeny follows (Rojas et al., 2016; Shi & Rabosky, 2015). Vertical black bars, from left to right, indicate: (1) Noctilionoidea, (2) Phyllostomidae, (3) and Stenodermatinae, respectively. The presence/absence of a protein product for S- and L-opsins was assayed by IHC on flat mounted retinas. The presence of protein is indicated by a filled color marker (OPN1SW – purple and OPN1LW – green), and its absence by a white marker. Finally, the OPN1SW protein assay for an additional six species (*Tadarida brasiliensis, Phyllostomus hastatus, Sturnira tildae, Sturnira ludovici, Platyrrhinus dorsalis* and *Chiroderma villosum*) represented by museum specimens failed and therefore, these species are not shown.

**Figure 3.**
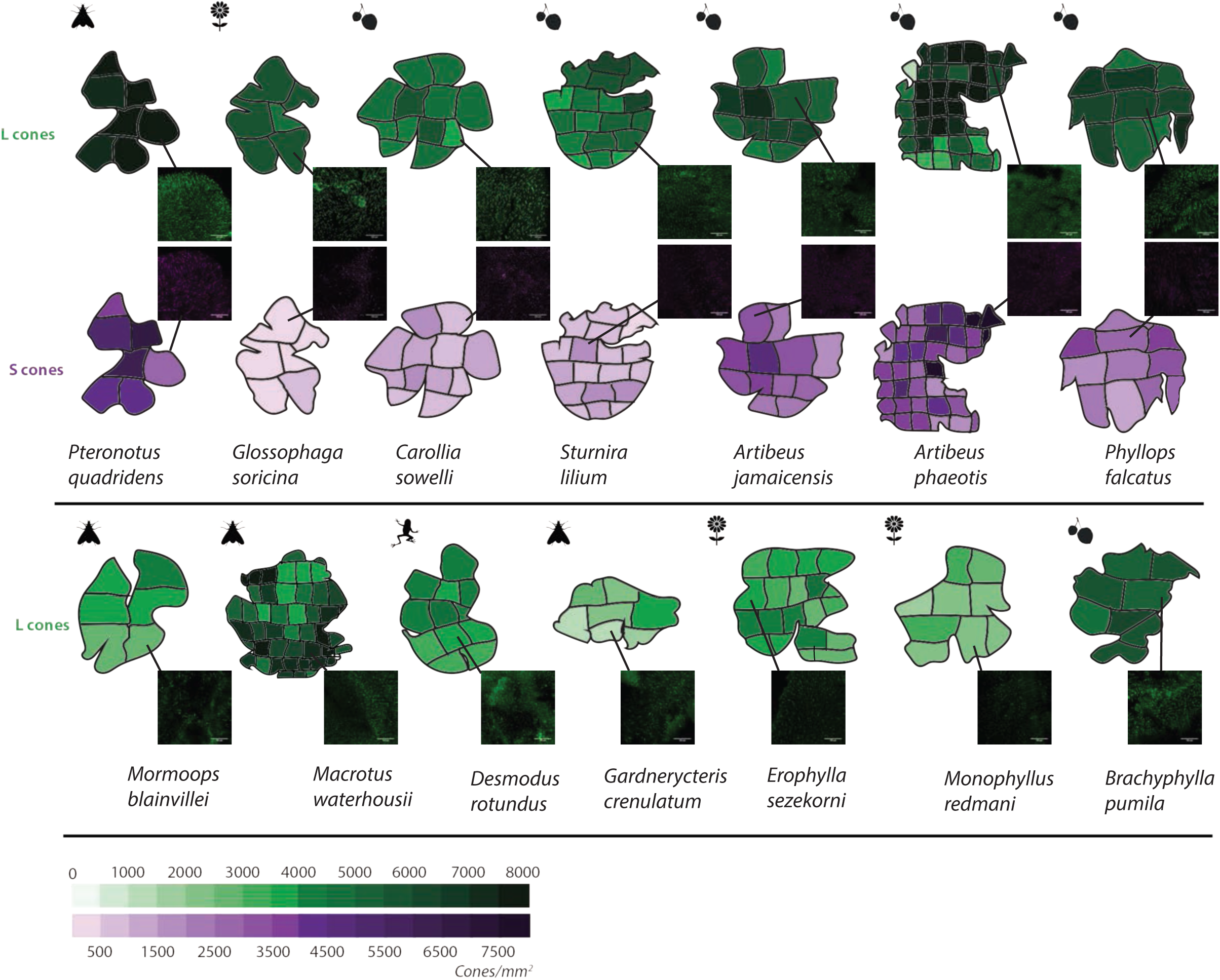
L and S opsin cone distribution in 14 representative noctilionoid vat species. Density maps of L and S opsin cone topography in 14 noctilionoid bat species. For each species, a representative dissected flat-mounted retina is shown. Insets are representative IHC magnifications of flat mounted retinas immune-stained for either L- or S-opsins in the highlighted region. Dietary types are indicated with the following symbols: invertebrates – moth, vertebrates – frog, fruit – fruit and nectar/pollen – flower. Measured opsin densities (0–8000 cones/mm^2^) are represented by the following color scales: L-opsins – green and S-opsins – purple.

### Mismatches between opsin transcript and protein repertoires

Comparison of *OPN1SW* transcript and protein data revealed several conflicts in species-specific absences. Seven lineages possessed *OPN1SW* transcripts but lacked the OPN1SW protein (Figure 4). Four of these species possessed apparently intact transcripts (*Molossus molossus*, *Tonatia saurophila*, *Gardnerycteris crenulatum*, *Monophyllus redmani*), while the remaining three included *P. parnellii* (an in-frame deletion), *E. bombifrons* (transcribed introns), and *C. brevicauda* (a premature stop codon). Among species lacking both transcript and protein, *OPN1SW* sequence data obtained via manual PCR or assembled genomes confirmed the open reading frame appears to be intact in *Chilonatalus micropus* and *Brachyphylla pumila*, but is disrupted in *Mormoops blainvillei*, *Macrotus waterhousii*, and *Desmodus rotundus.*

**Figure 4.**
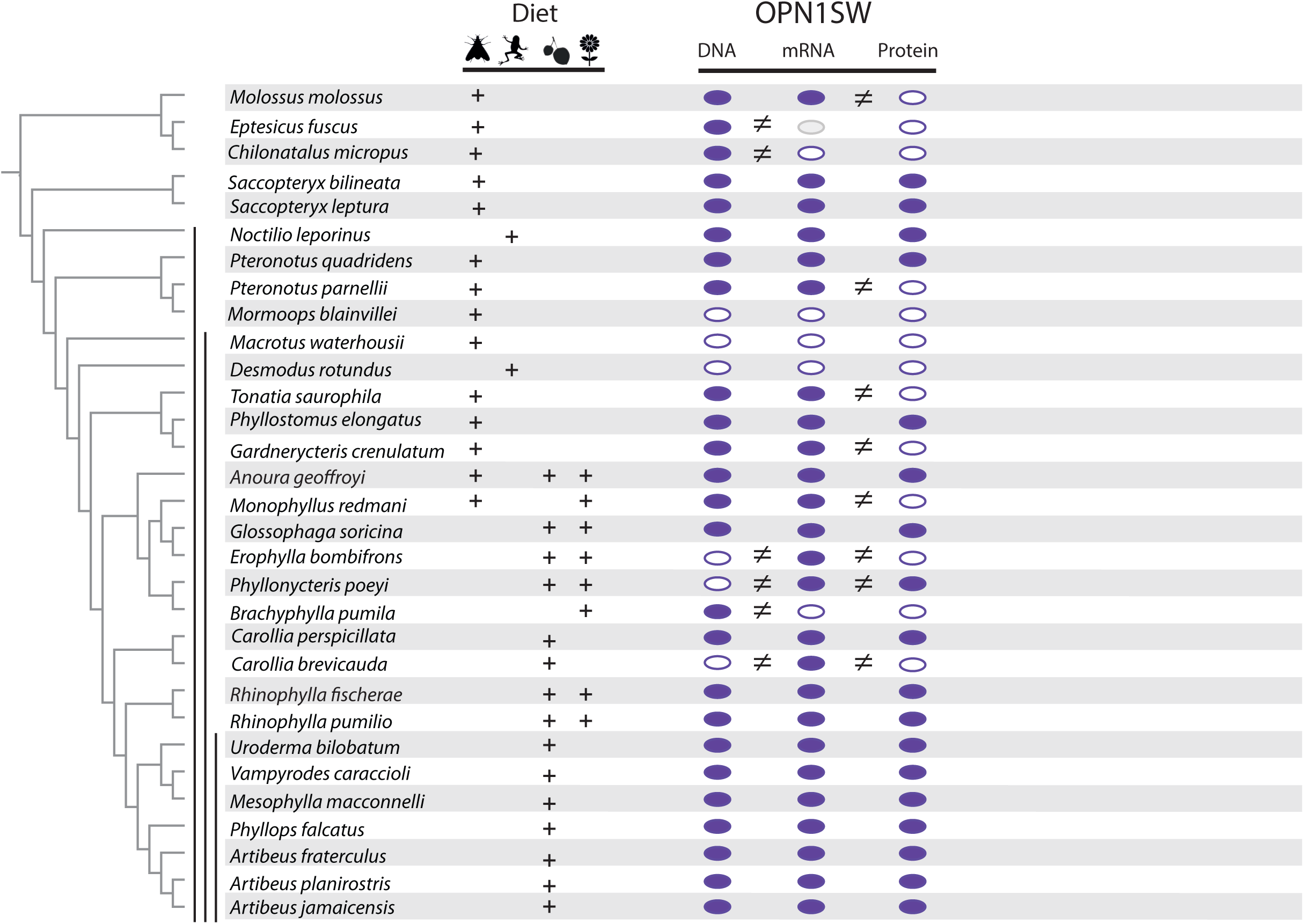
Distribution of an intact open reading frame (ORF), mRNA transcript and protein for the OPN1SW photopigment in ecologically diverse noctilionoid bats. The composition of species diet follows Rojas et al. (2018), dietary types are indicated with the following symbols: invertebrates – moth, vertebrates – frog, fruit – fruit and nectar/pollen – flower. The species phylogeny follows (Rojas et al., 2016; Shi & Rabosky, 2015). Vertical black bars, from left to right, indicate: (1) Noctilionoidea, (2) Phyllostomidae, (3) and Stenodermatinae, respectively. RNA-Seq data was generated to both infer the presence of an intact ORF (in combination with genomic and PCR sequence data) and determine the presence of an expressed mRNA transcript. The presence/absence of a protein product for S-opsins was assayed by IHC on flat mounted retinas. The presence of an intact ORF, mRNA and protein are indicated by a filled color marker, and its absence by a white marker. Missing data (i.e. species for which we were unable to obtain tissue) are indicated with a grey marker. Mismatches between intact ORFs and transcripts, or between transcripts and protein data are indicated by an inequality symbol. Note: OPN1SW protein assays for *P. quadridens* revealed polymorphisms within the sample, and we recorded positive OPN1SW assays in some *P. poeyi* individuals despite an apparent disrupted ORF.

In species in which a protein was detected, we also found a corresponding transcript. However, in the case of *P. poeyi*, data from four individuals showed S-cones in all, despite indels and retained introns detected in the transcript of a fifth different individual (confirmed via PCR). In contrast to the mismatches between *OPN1SW* mRNA and OPN1SW protein, we found complete correlation between the presence of the transcript and protein for OPN1LW.

### Molecular evolution of opsin genes

To compare and contrast the evolution of opsins, tests of divergent selection modes were performed for each of the three opsin genes (*OPN1SW*, *OPN1LW* and *RHO*) for three types of lineages: (1) those with S-cones; (2) those without S-cones but with *OPN1SW* transcripts and an intact *OPN1SW* open reading frame; and (3) those without either S-cones or *OPN1SW* transcripts (see Figure supplement 2). There was a significantly higher *ω* in lineages with a pseudogenized. *OPN1SW* in both the *OPN1SW* gene (*ω*_background_ = 0.11; *ω*_OPN1SW.intact_ 0.07; *ω*_OPN1SW.pseudo_ 0.79; *χ*^2^_(2)_ = 129.9, *P* < 1.0e-5), and the *OPN1LW* gene (*ω*_background_ = 0.08; *ω*_OPN1SW1.intact_ = 0.08; *ω*_OPN1SW.pseudo_ = 0.17; *χ*^2^_(2)_ = 6.9, *P* = 0.03). In contrast, we found no differences in *ω* across the different lineages for *RHO*, indicating similarly strong, negative selection across all lineages (Table S2). To test the influence of diet on rates of molecular evolution, we compared *ω* for all three opsin genes between frugivorous and non-frugivorous lineages. There were no differences in rates for *OPN1SW* or *OPN1LW*, but background branches (non-frugivore species) had a significant and slightly higher *ω* for *RHO* (*ω*_background_ = 0.04; *ω*_fugivory_ = 0.02; *χ*^2^_(1)_ = 110, *P* = 9.1e-i 4; Table S3).

### Ecological correlates of opsin presence and density

We compared the locations and densities of long-wavelength-sensitive cones (or L-cones expressing OPN1LW protein) and short-wavelength-sensitive (or S-cones expressing OPN1SW protein) in the whole, flat-mounted retinas of adult bats for 14 species for which we had sufficient specimen replicates (Table S4). We found photoreceptor density varied among examined species (Figure 3, Table S4), with mean cone densities ranging from 2,500 to 7,500 cones/mm^2^ for L-cones, and 327 to 5,747 cones/mm^2^ for S-cones (Figure 3, Table S4). In every case, the density of S-cones was lower than that of L-cones. Densities of both cone types tended to be highest near the center of the retina in all species. Bat species with S-cones had significantly higher densities of L-cones, with the presence of S-cones increasing L-cones by 43% in the natural logarithm scale or 154% in the linear scale, explaining on average ∼24% of the variance in density between species (Table 1).

**Table 1.** Summary of Bayesian regression models of the presence of S-cones or the density of L-cones as a function of predictor variables. Posterior estimates are shown as medians, and 2.5%, 97.5% high probability density intervals (in parentheses).

| Modeled response | Predictor | Coefficient | $R^2$ |
| --- | --- | --- | --- |
| S-cones (presence/absence) | Fruit in diet (prevalence/non) | 3.63 (1.82, 6.72) | 0.58 (0.18, 0.83) |
| ln(L-cones) | Fruit in diet (prevalence/non) | 0.22 (-0.21, 0.66) | 0.00 (0.00, 0.30) |
| ln(L-cones) | S-cones (presence/absence) | 0.43 (0.09, 0.78) | 0.24 (0.00, 0.51) |

We tested the relationship between the presence of S-cones and diet using Bayesian hierarchical models with the phylogenetic structure of the data as a species-specific effect. Comparisons of the Deviance Information Criterion (DIC), for which lower values indicate better fit of the model to the data, showed that the best-fitting model included frugivory as an explanatory variable (Figure supplement 4). The predominance of fruit in the diet increases the odds of having S-cones roughly 38 times, and explains about 58% of the between-species variance in the presence of the S-cone (Table S5, Figure 5).

**Figure 5.**
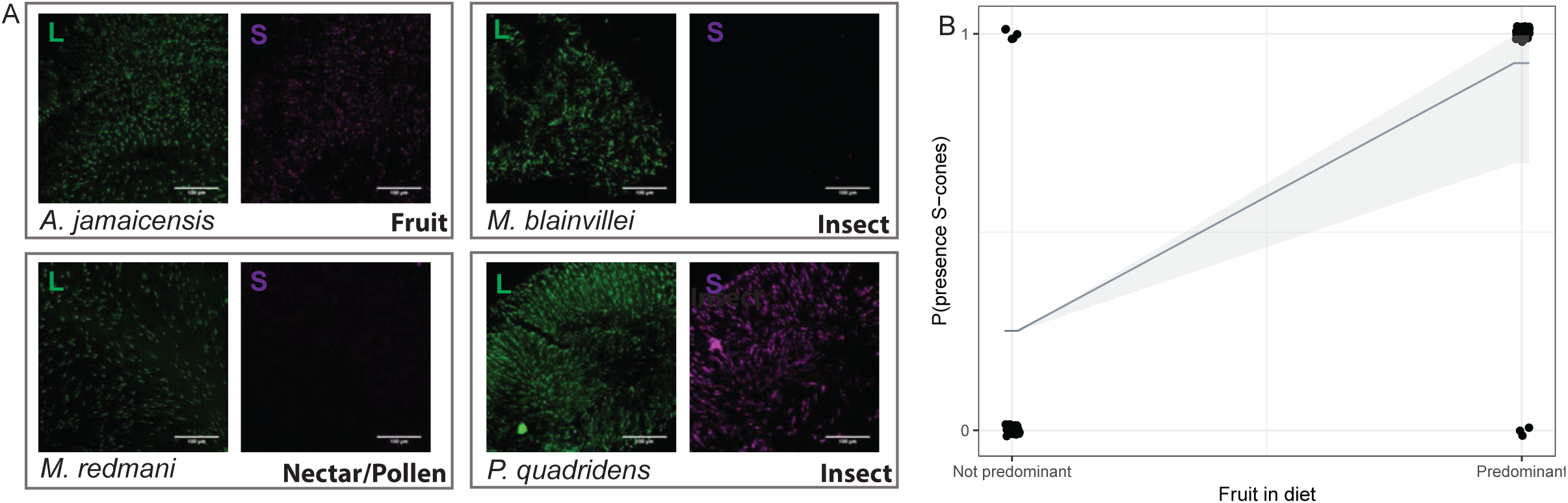
S-opsin presence is correlated with diet. A: Representative IHC magnifications of flat mounted retinas immune-stained for both L and S opsin for four species representatives of the diversity of phenotypes observed. Fruit-based diet: *Artibeus jamaicensis*, pollen/nectar-based diet: *Monophyllus redmani*, and insect-based diet: *Mormoops blainvillei* and *Pteronotus quadridens.* B: Modeled probability of the presence of S-cones as a function of the predominance of fruit in diet. The posterior distributions of parameters for this model are summarized in Table 1.

## Discussion

Studies of mammalian photopigment genes have provided important insights into molecular adaptations underpinning vision, but typically have assumed correspondence between open reading frames and the protein phenotypes. By studying opsin gene sequences, transcripts and proteins across a radiation of New World bats (suborder Yangochiroptera, superfamily Noctilionoidea), we discovered unexpected diversity in genotypes and phenotypes that points to hitherto unsuspected differences in visual perception among species. Within-lineage mismatches between DNA, mRNA, and protein also revealed multiple molecular mechanisms underlying similar visual phenotypes, suggesting various stages in the loss of color vision.

In the case of *opsin 1*, *short wave sensitive* (*OPN1SW*) gene, while the presence (or absence) of the transcript and protein was consistent across most species, there were also multiple exceptions. In contrast, for the *opsin 1*, *long wave sensitive* (*OPN1LW*) gene, both the transcript and protein were detected in all taxa examined. Additionally, on the basis of concordance between DNA and RNA, and in line with other studies [e.g. (Zhao, Ru, et al., 2009)], we confirmed that rhodopsin was retained and appears to be expressed in all taxa.

We are able to infer divergent evolutionary histories of diverse bat lineages by mapping the presence of an intact *OPN1SW* coding sequence, mRNA and protein onto a phylogeny spanning seven bat families, and combining this with molecular evolution analyses. While most species examined appeared to possess functional OPN1SW proteins, we were also able to identify multiple routes of functional degradation in those that did not. The relationship we observed between protein phenotype and corresponding ecology suggests frugivory, a novel ecological niche that evolved multiple times in the radiation of noctilionoid bats, was a causal factor shaping differences in S-cone retention.

We identified three separate routes by which S-opsin cones fail to form (Figure 4 and 6). First, we found evidence of pseudogenization arising from indels and/or stop codons, such as in *Desmodus rotundus, Macrotus waterhousii*, and *Mormoops blainvillei* (Figure 6B). Second, in the one sampled representative of the family Natalidae, *Chilonatalus micropus*, we found evidence of an intact gene sequence, yet the transcript was not detected, as was also the case in *Brachyphylla pumila* (Figure 6C). Finally, we recovered *OPN1SW* transcripts but were unable to detect the corresponding S-cone protein in a number of taxa (Figure 6D): *Molossus molossus*, *Pteronotus pamellii*, *Tonatia saurophila*, *Gardnerycteris crenulatum*, *Monophyllus redmani*, *Erophylla bombifrons*, and *Carollia brevicauda*. Of these, we found the open reading frame was intact in all but the last two species.

**Figure 6.**
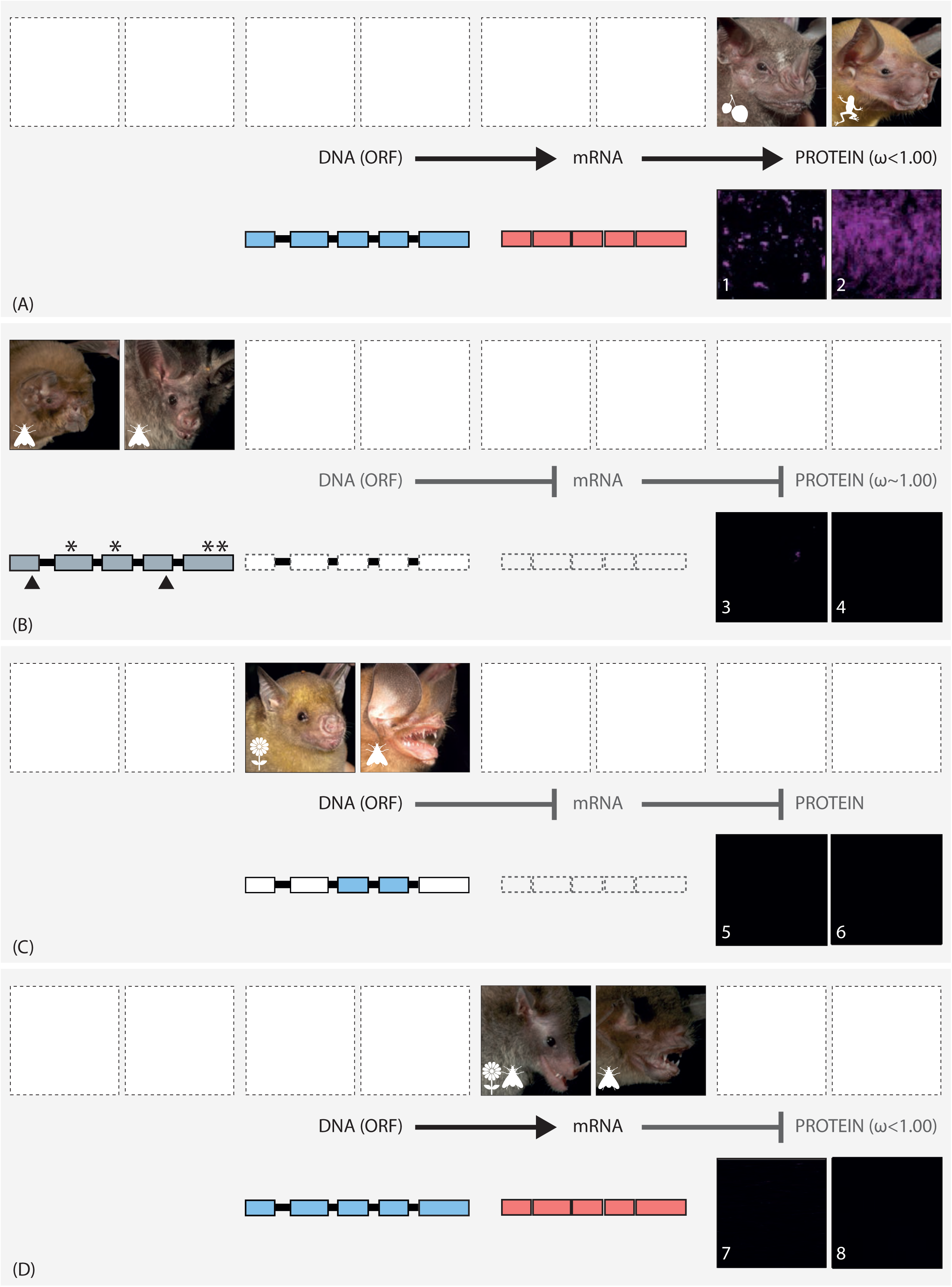
The putative routes explaining variation in S-cone presence in noctilionoid bats. In each panel, upper images (left and right) show the gross phenotype of the eye in representative bat species, and lower images (numbered) show IHC magnifications of their respective flat mounted retinas immune-stained for S-opsin. Diets are depicted with the following symbols: invertebrates – moth, vertebrates – frog, fruit – fruit and nectar/pollen – flower. A. Information in the intact DNA Open Reading Frame (blue) is transcribed to form mRNA (red), which is then translated into OPN1SW. Codon analyses reveal purifying selection. Example species: (L + 1) *Artibeus jamaicensis*; (R + 2) *Noctilio leporinus.*
B. The DNA ORF is disrupted (grey) by the presence of STOP codons (^∗^) and in-dels (black triangles). Neither *OPN1SW* mRNA (dashed boxes) nor OPN1SW is detected. Codon analyses reveal relaxed selection. Example species: (L + 3) *Mormoops blainvillei*; (R + 4) *Macrotus waterhousii*.
C. Although the DNA ORF (blue) appears to be intact, information is not transcribed to mRNA (dashed boxes), and no OPN1SW is detected. Example species: (L + 5) *Brachyphyllapumila* (R + 6) *Chilonatalus micropus.*
D. Information in the intact DNA ORF (blue) is transcribed to form mRNA (red), however, the OPN1SW is not detected. Codon analyses reveal purifying selection. Example species: (L + 7) *Monophyllus redmani* (R + 8) *Pteronotus parnellii.*

Pseudogenization, whereby a non-essential gene loses some functionality, occurs relatively frequently within mammalian genomes, with previous estimates suggesting several thousand pseudogenes may be present per genome [e.g. (Torrents et al., 2003)]. Instances of disruption in reading frames of visual cone-opsins for *OPN1SW* and, to a lesser extent, *OPN1LW*, are not unusual, and have been previously reported in mammals multiple times [e.g. (Emerling et al., 2015; Emerling et al., 2017; Zhao, Rossiter, et al., 2009)]. However within bats, previous studies suggested the *OPN1SW* gene had only been lost in certain lineages of the suborder Yinpterochiroptera [e.g. (Emerling et al., 2015; Kim et al., 2008; Müller et al., 2007; Zhao, Rossiter, et al., 2009)], and thus the OPN1SW protein, and the associated putative UV-sensitive vision, were thought to be prevalent across Yangochiroptera [e.g. (Emerling et al., 2015; Gorresen et al., 2015; Müller et al., 2009; Winter et al., 2003)]. By revealing pervasive losses of S-cones in members of the Yangochiroptera, our findings both upend current understanding of visual perception across suborders of bats, and complicate hypotheses relating the evolution of specialized sensory behaviors to changes in opsin repertoires [e.g., (Zhao, Rossiter, et al., 2009)]. Additionally, in all species for which we were able to obtain the 5’ region of the open reading frame, we found that the putative canonical start codon was located 9-bp downstream from that previously annotated in bats and other mammals, such as humans. This has previously been described in some rodents [e.g., (Shimmin et al., 1997; Zerbino et al., 2018)], further highlighting the opsin peptide divergence found within the bat clade.

In the second inferred form of loss, the open reading frame is seemingly intact but appears not to be transcribed. While only manual PCRs of ∼100 codons of the *OPN1SW* gene from *Chilonatalus micropus* and *Brachyphylla pumila* support this type of loss, a previously published sequence confirms the presence of a functional open reading frame in an additional species from the same family as *Chilonatalus* (Emerling et al., 2015). As such mismatches between DNA and mRNA have not previously been documented in bat opsins, more work is needed to both explore the extent to which this occurs, as well as the molecular mechanism that underlies it.

Perhaps the most unusual result of this study was the apparent failure of an intact transcript to be translated into protein in several taxa. Although the underlying cause of these disruptions in protein translation can at this point only be inferred, close inspection of the transcripts involved suggests that more than one molecular mechanism may be involved. In two taxa, *Erophylla bombifrons* and *Phyllonycteris poeyi*, examination of the assembled transcript contained retained introns (confirmed via BLAST). Transcriptional readthrough is well documented [e.g., (Gaidatzis et al., 2015; Vilborg & Steitz, 2017)] and such carryover of intronic sequence into OPN1SW would interfere with protein formation. The presence of introns in *OPN1SW* mRNA has previously been documented in the blind mole-rat *Spalax ehrenbergi*, which also lacks S-cones (David-Gray et al., 2002; Esquiva et al., 2016). In another case, multiple individuals of *Pteronotus quadridens* showed variation in S-cone presence among conspecifics, indicating possible ongoing degradation of protein synthesis. These mechanisms are not mutually exclusive: a similar result was found for *P. poeyi*, for which S-cones were detected in four animals, alongside a seemingly non-functional transcript detected in another that contained retained intronic sequences as well as a 4 base-pair insertion in exon 3, again verified by manual PCR.

Taken together, our findings reveal assessments of visual perception based purely on genotypic analyses of opsin sequences, or RNA transcripts, can be misleading, and may even obscure the evolutionary processes and ecological agents of selection. Although variation in the complement of photoreceptors across vertebrates is usually explained by disruptions to the protein-coding sequence [e.g., (Mundy et al., 2016; Zhao, Rossiter, et al., 2009)], our findings of mismatches between genotype and phenotype also indicate a role for transcriptional and even translational control in this process.

Our results further allow us to hypothesize the developmental processes contributing to variation in visual perception among bat taxa. Unlike species of *Mus* (Ortín-Martínez et al., 2014), in which many cones express more than one photoreceptor, almost all cones in the bats we analyzed expressed a single opsin type (OPN1SW or OPN1LW). Given this constraint, if S-opsin cones were lost through developmental conversion into L-opsin cones, then, species with S-opsin cones should have fewer L-opsin cones than species without S-opsin cones. Instead, species with S-opsin cones actually have more L-opsin cones. Therefore, our findings are more consistent with the loss of S-opsin cones in Yangochiroptera occurring during the process of cone determination. Comparisons of photoreceptor development across ontogeny are needed to test this hypothesis.

Considering the distribution of losses throughout the phylogeny, and the many, diverse forms of loss we identified, independent losses of S-cones must have occurred during the radiation of noctilionoid bats. At a much broader phylogenetic scale, independent losses of S-opsin cones have been documented more than 20 times throughout mammalian evolution (Emerling et al., 2015; Hunt & Peichl, 2014), and monochromatic —often nocturnal— species are usually closely related to dichromatic relatives. Cases of loss-of-function in *OPN1SW* genes caused by the accumulation of premature stop codons and other mutations have been documented across many vertebrate lineages, and are generally inferred to require many generations (Kawamura & Kubotera, 2004). In contrast, decreases in gene expression, which may represent the first step toward functional loss, can evolve rapidly within populations (e.g., in cave fish) (Tobler et al., 2010).

Mismatches between mRNA transcripts and protein assays for OPN1SW in bats reveal the evolution of photoreceptor composition and repartition to be flexible compared to pseudogenization. *OPN1SW* genes with intact open reading frames in lineages that lack the OPN1SW protein show few amino acid substitutions. This suggests mismatches between transcription and expression evolve rapidly and may explain low rates of evolution whether or not the S-cones are present (Table S2). Once the *OPN1SW* gene is pseudogenized, however, the degradation of the gene seemingly becomes irreversible and a manifold increase in the protein-coding substitution rate ensues. Both frugivorous and non-frugivorous lineages experience strong purifying selection to conserve ancestral function, highlighting post-transcriptional regulation as a more direct response to ecology than pseudogenization of the relevant opsin (Table S4). These results therefore suggest that protein composition should more closely reflect visual ecology than high rates of sequence evolution and pseudogenization in the relevant opsin, as the latter only responds to long-term functional loss.

Despite these observed widespread losses, the conservation of S-cones, *OPN1SW* transcription, and protein-coding sequences in most species implies that S-cones have an important function in most noctilionoid bats. To identify the likely ecological determinants of variation in S-cone presence across the clade, we applied phylogenetic regressions, and found the predominance of fruit consumption —and notably not of plant or flower-visiting, or insectivory— was the single most powerful explanatory factor. This result strengthens the role of diet as the primary agent of selection that maintains S-opsin function, and counters previous speculation that nectarivorous New World leaf-nosed bat species rely on UV reflectance to locate flowers through short-wave vision (Winter et al., 2003). Instead, we found some predominantly nectarivorous species such as *Erophylla spp.* and *Lonchophylla robusta* have lost their S-cones, while others have retained them (e.g., *Anoura geoffroyi*). This both suggests more than one strategy to finding flowers has evolved among New World leaf-nosed bats, and contrasts with members of the primarily frugivorous subfamily Stenodermatinae, which invariably retain their S-cones (Figure 2).

In primates, the ability to discriminate between fruit and foliage has been identified as an ecological advantage of daylight trichromacy (Fedigan et al., 2014), and bats that include fruit in their diets could benefit from some degree of color discrimination. Although bats that possess both L- and S-cones are either monochromatic or dichromatic in daylight, and conditionally dichromatic or trichromatic in mesopic (i.e., crepuscular) conditions (Zele & Cao, 2015), the ability to detect UV light might be important in fruit detection. Indeed, several stenodermatine bat species analyzed here appear to be able to perceive UV light based on inferences from spectral tuning of OPN1SW photopigments (Zhao, Rossiter, et al., 2009), behavioral tests of responsiveness to UV light (wavelengths < 400 nm) (Gorresen et al., 2015; Winter et al., 2003), and electroretinographic recordings of visual sensitivity (Müller et al., 2009). In plant species with animal-mediated seed dispersal, dispersers are attracted to ripening fruits via a combination of visual and olfactory cues that include the emission of attractive volatile compounds, a reduction of toxic compounds, and changes in texture and color [e.g., (Goff & Klee, 2006; Weiblen et al., 2010; Wendeln et al., 2000)]. Increases in UV reflectance also correlates with the decline of chlorophyll and ripening in several fruits [for reviews see (Blanke & Lenz, 1989; Lagorio et al., 2015; Thies et al., 1998)]. Given that UV reflectance has been documented in paleotropical fig species (Lomascolo et al., 2010), selection for UV perception at twilight might explain the retention of S-cones in primarily frugivorous lineages. Indeed, we speculate the persistence of UV vision in the ancestor of the stenodermatine bats may have facilitated their invasion of a fig-dominated dietary niche, contributing to their rapid taxonomic and ecological diversification (Rojas et al., 2018).

Since the consumption of fruit arose as an evolutionary innovation within Yangochiroptera, selection for this novel niche cannot explain the ancestral or present-day persistence of S-cones in non-frugivorous species (Figure 2). While the retention of S-cones in bats that feed on animals (e.g., *Chrotopterus auritus* and *Noctilio leporinus*) is surprising, recorded polymorphisms in the presence of S-cones among some individuals of insectivorous *P. quadridens* could point to an ongoing loss of S-cones in some of these lineages. Comparisons with L-opsin cones offer insights into the general functional role of S-cones. While fruit consumption was not a covariate of L-opsin cone density, the presence of S-cones is associated with an estimated 43% increase in the density of L-opsin cones (Table 1), consistent with both types of cones serving a common functional role. This role could be to increase visual sensitivity at dusk, similar to crepuscular and occasionally diurnal Old-World fruit bats (Zhao, Rossiter, et al., 2009). As a result, light capture, instead of the detection of novel visual cues, might have been the primary agent of selection on the density of S- and L-opsin cones in non-frugivorous lineages, as well as ancestral bats. The presence of S-cones in ancestral noctilionoid bats might have then allowed them to take advantage of fruit as a novel fruit source during their subsequent diversification, in a case of trait exaptation.

## Conclusions

Vertebrate sensory adaptations have been the focus of intense research over the last fifteen years, and bat vision has not been an exception [e.g., (Eklof & Jones, 2003; Feller et al., 2009)]. These studies have upended formerly prevalent views on bat biology and behavior, finding increasingly important roles for vision (and olfaction) in orientation and food localization, even among echolocating species [e.g., (Altringham & Fenton, 2003; Korine & Kalko, 2005)]. But while previous analyses of isolated lineages revealed the existence and extent of mesopic and photopic cone-based vision in fewer than five noctilionoid species (Müller et al., 2009; Winter et al., 2003), we examined dozens of ecologically diverse bat lineages throughout the Yangochiroptera suborder and found a much more complex picture than was previously assumed.

We uncovered mismatches between *OPN1SW* transcripts and S-opsin cone phenotypes, as well as widespread S-cone loss, with conservation linked to multiple factors including a novel dietary niche. Our results also highlight the importance of rapid trait loss in evolution, with apparent shifts in translation that precede pseudogenizing changes in open reading frames. As genotype-centered analyses would miss important functional changes, our study illustrates the importance of a broad comparative approach when studying sensory innovation, loss, and adaptation, as well as their influence on taxonomic and ecological diversification.

## Materials and methods

### Species sampling and tissue preparation

We obtained eye tissue from 59 New World bat species, of which 49 were collected from the wild and 34 from museums, with 24 species common to both sources (Table S1). All wild bats were captured with traps set in forests and/or at cave entrances, were handled, and then euthanized by isoflurane overdose, under appropriate research and ethical permits (see Supplementary Information).

### RNA sequence analysis

Intact eyes were placed in RNAlater and incubated at 4°C overnight and then frozen. Total RNA was isolated using Qiagen RNeasy Mini kits with the addition of DTT and homogenization using a Qiagen TissueLyser. Following QC, total RNA from each individual was used to construct a cDNA library using the Illumina TruSeq^®^ RNA v2 kit. Pooled libraries were sequenced (NextSeq 500). Eye transcriptomes were generated for 45 individuals (38 species) including biological replicates of *Pteronotus parnellii* (*n* = 4), *Artibeus jamaicensis* (*n* = 4) and *Phyllops falcatus* (*n* = 2). Raw reads were trimmed, and clean reads were assembled with Trinity v.2.2.0 (Grabherr et al., 2011) (see Supplementary Information).

We tested for the presence of the three focal gene transcripts (*RHO*, *OPN1SW* and *OPN1LW*) in each bat transcriptome using a reciprocal best hit blast approach against the full set (n=22,285) of human protein-coding genes from Ensembl 86 (Yates et al., 2016). To confirm the absence of *OPN1SW* sequence, we performed additional steps in several species. First, we cut, *trans*-chimeras, which can prevent detection by reciprocal blast (Yang & Smith, 2013), and repeated the reciprocal blast. Second, we manually screened sequences that were initially identified as matching *OPN1SW*, but did not pass initial blast filtering (see Supplementary Information). Recovered opsin gene sequences have been submitted to GenBank (accession numbers XXX-XXX).

### Immunohistochemistry (IHC) and photoreceptor quantification

Eyes were fixed overnight at 4°C in PBS 4% PFA, and then stored in 1% PFA until processing. Museum ethanol-fixed specimens were first rehydrated through PBS-methanol series. Retinas were dissected from the eyeballs, oriented and flattened by making radial incisions (see Supplementary Information). IHC was then performed on dissected retinas following standard procedures as in Ortin-Martinez *et al.* (2014), with the appropriate mixture of primary and secondary antibodies (see Supplementary Information for details). For each species, IHC was performed on replicates from at least three individual retinas. We considered no labeling to indicate a true loss of the cone type following references (Müller et al., 2009; Müller et al., 2007; Ortín-Martínez et al., 2014). Labelled retinas were then imaged on a confocal microscope at 20X (LSM710; Zeiss Microscopy, see Supplementary Information for details). The entire retina surface was captured using 512×512 pixel tiles and counted using a 3D object counter plugin using Fiji (ImageJ). The accuracy of this approach was verified manually (Supplementary Information). Opsin density for each species was calculated by averaging across three individuals. We also characterized the spatial distribution of L- and S-cone densities for 14 species.

### Opsin gene evolution

We used aligned sequences from the transcriptomes of 38 species together with those from six noctilionoid genomes (Zepeda Mendoza et al., 2018) to estimate rates of molecular evolution of visual opsin genes (*OPN1SW*, *OPN1LW*, and *RHO*) in focal bats. First, we tested for divergent selection modes among species that had S-opsin cones, lacked the S-opsin cones but had an intact mRNA sequence, and those that lacked the S-opsin cones but either did not have *OPN1SW* transcripts or had a pseudogenized *OPN1SW* sequence (Figure supplement 2) using the Branch Model 2 of codeml in PAML 4.8a (Yang, 2007). Second, we applied the same approach to test divergent selection modes between frugivorous and non-frugivorous bat species (Figure supplement 2; gene alignments have been submitted to DRYAD XXXX).

### Ecological correlates of cone presence and density

To determine whether cone phenotypes are explained by dietary specialization, we applied the hierarchical Bayesian approach implemented in the R packages MCMCglmm and mulTree (Guillerme & Healy, 2014; Hadfield, 2010), using a random sample of trees from the posterior distribution of published phylogenies (Rojas et al., 2016; Shi & Rabosky, 2015). We modeled S-cone presence as function of diet, and compared models using the Deviance Information Criterion (DIC) (Gelman, 2004). Using the predictor variable from the best-fit model for presence/absence of S-opsin cones, we then repeated this approach to explain L-cone density (see Supplementary Information).

## Acknowledgments

We thank M. Agudo-Barriuso, P. K. Ahnelt, L. Peichl, G. Tsagkogeorga and staff at the Queen Mary Genome Centre for advice, lab assistance and protocols. For help with permits and field support in the Dominican Republic, we thank J. Almonthe, M.E. Lauterbur, Y.M. León, M. Nuñez, and J. Salazar; in Peru, F. Cornejo, J. Pacheco, J. Potter, H. Portocarrero, M.K. Ramos, E. Rengifo, J.N. Ruiz, C. Tello; and in Puerto Rico, A. Rodriguez-Duran, and N. Ann. For access to museum specimens, we thank N. Simmons (American Museum of Natural History). For help with permits and field support in Belize, we thank M. Howells and the Lamanai outpost lodge staff, N. Simmons and B. Fenton. For providing bat images we thank E. Clare. Results in this paper were obtained using the high-performance LI-RED computing system at the Institute for Advanced Computational Science at Stony Brook University, and Queen Mary’s MidPlus computational facilities supported by QMUL Research-IT and funded by EPSRC grant EP/K000128/1. L.M.D. and S.J.R. were supported in part by the National Science Foundation DEB-1442142, K.E.S. by DEB-1442314, and E.R.D. by DEB-1442278. Additionally, S.J.R and K.T.J.D. were supported by the European Research Council (ERC Starting grant 310482 [EVOGENO]). This research was conducted under research permits VAPB-01436 in the Dominican Republic, and 0002287 in Peru.

## Author contributions

AS, KTJD, ERD, LMD, SJR and KES designed research; AS, KTJD, KY, LRY, PD and LMD performed research; AS, KTJD, LMD, SJR, KES and LRY contributed new samples, reagents or analytic tools; AS, KTJD, LRY and LMD analyzed data; AS, KTJD, LRY, BPH, ERD, LMD, SJR and KES wrote the paper.

## Supporting Information

### Supplementary materials and methods

#### Species sampling

We obtained eye tissue samples from a total of 263 eyes from 232 individuals, representing 59 bat species from seven families. This includes three families from the focal Noctilionoidea superfamily and four closely related outgroup families. Specimens used for this study were either wild-caught animals or obtained from museum collections (see Table S1 for full species and permit information). Wild-caught specimens from 49 bat species were collected following the approved IACUC protocols and site-specific permits. Bats were sampled from wild populations, and caught using mist nests set in forests and/or at cave entrances. Animals were handled following IACUC and site-specific protocols to minimize stress and were euthanized using an excess of isoflurane. Fresh bat specimens used for RNA-Seq analyses (*n*_*RNA-Seq*_ = 45) were sampled in the Dominican Republic and Peru, and those for immunohistochemistry were sampled in the Dominican Republic, Puerto Rico, Belize, and Trinidad. Additional eye samples from 34 species were dissected from specimens from the American Museum of Natural History (AMNH). For immunohistochemistry, five bat species had replicates that were both wild-caught and from museum collections and exhibited the same phenotype, highlighting the robustness of the experiments. Due to preservation methods, the AMNH samples were only suitable for immunohistochemistry (IHC) and so were not included in the transcriptomic study (see Table S1 for AMNH specimen identification numbers).

#### RNA sequence analysis

Shortly after death, intact eyes were excised and placed in RNAlater and incubated at 4°C overnight before being stored at −180°C in vapor-phase liquid nitrogen, or at − 80°C in a freezer. Frozen eye tissues were placed in Buffer RLT with added Dithiothreitol (DTT) then homogenized using a Qiagen TissueLyser. Total RNA was then extracted using Qiagen RNeasy^®^ Mini kits following the manufacturer’s protocol. Following extraction, RNA integrity was assessed using an Agilent 2100 Bioanalyzer and RNA concentration was measured using a Qubit Fluorometer. Library preparation was performed using Illumina TruSeq^®^ RNA Sample Preparation v2, with 500 ng of total RNA used for each sample. Constructed libraries were pooled and sequenced on the NextSeq500 High Output Run (150 cycles) to give 2×75 base-pair (bp) paired-end (PE) reads at the Genome Centre, Queen Mary University of London. Using the above approach, we sequenced eye transcriptomes from 45 individuals, representing 38 bat species; including biological replicates for three species (*Pteronotusparnellii: n* = 4; *Artibeus jamaicensis*: *n* = 4 and *Phyllops falcatus: n* = 2).

We assessed the quality of the short-read data with FastQC v.0.11.5 (http://www.bioinformatics.bbsrc.ac.uk/projects/fastqc). Raw reads were trimmed with Trimmomatic-0.35 (Bolger et al., 2014), with the following settings LEADING.3 TRAILING.3 SLIDINGWINDOW.4.15 MINLEN.36. Cleaned reads were assembled into *de novo* transcriptomes with Trinity v.2.2.0 (Grabherr et al., 2011), using default parameters.

#### Opsin gene annotation and examination of CDS

We used a combination of transcriptomic and genomic data to establish if the coding sequences (CDSs) of the three focal genes (*OPN1SW*, *OPN1LW* and *RHO*) were intact, and whether or not the mRNA of these genes was expressed in the eyes of the bats under study. We used a reciprocal best hit blast approach to establish whether or not transcripts corresponding to the three visual pigments (*OPN1SW*, *OPN1LW*and *RHO*) were present in each of the assembled transcriptomes. Sequences representing the protein products encoded by 22,285 human protein coding genes were downloaded from Ensembl 86 (Yates et al., 2016), for each gene product only the longest protein sequence was retained. These sequences were then used as tblastn (blast+ v.2.2.29) queries against each of the 45 bat transcriptome databases, the top hit was kept with an e-value cut-off <1e^−6^. Reciprocal blasts were then carried out using blastx (blast+ v.2.2.29), with bat transcripts as queries against the human protein database, again only keeping the top hit and e-value <1e^−6^. Percentage coverage of each bat hit against the human protein was calculated with the perl script analyze_blastPlus_topHit_coverage.pl available in Trinity utils. Candidate coding sequences (CDSs) were then extracted from the transcriptome assembly based on the blast coordinates using a custom perl script.

For transcriptome assemblies in which the *OPN1SW* sequence was not initially recovered, we undertook a number of additional steps to confirm that the sequence was not present. Firstly, as *de novo* transcriptome assemblies can create erroneous chimeric transcripts that may affect reciprocal blast results we followed the approach of (Yang & Smith, 2013) to reduce the number of transself-chimeras. Briefly, this involves performing a blastx search of the bat queries against the human protein database with an e-value cutoff of 0.01. Hits that met the default parameters of identity ≥30% and length ≥100 base-pairs are used to then search for either self-chimeras or multigene chimeras. Detected putative chimeras are then cut into segments, and retained if >100 base-pairs. This approach is only able to screen for *trans*-chimeras. Following chimera detection, the initial reciprocal blast described above was repeated. Lastly, we manually re-blasted all sequences that were initially identified as matching *OPN1SW*, but did then not pass the stringent reciprocal blast procedure.

Finally, we mapped the reads of *Carollia brevicauda* against *Carollia perspicillata* to validate the presence of a stop codon. Mapping was performed using the script align_and_estimate_abundance.pl available from Trinity utils, and RSEM v. 1.2.31 and BOWTIE v.1.1.2.

#### Verification of OPN1SW sequence with PCR

For three species we amplified the genomic region spanning exons 3–4 of *OPN1SW*, from DNA extracted from the same individuals used to generate the RNA-Seq data. DNA extractions were carried out using Qiagen DNeasy Blood & Tissue Kit (69504). We used previously published primers (Zhao et al., 2009). PCR mixtures consisted of 12.5 μl EconoTAQ Master Mix 2×, 3 μl of each primer (10 μM), 4 μl of genomic DNA (> 100 ng). On an Eppendorf Mastercycler ProS, PCR was carried out with a single cycle at 94°C for 2 min followed by 35 cycles of 94°C for 30 sec, annealing temperature of 55°C for 40 sec, 72°C for 1 min 30 sec, and finally a single cycle of 72°C for 10 min. PCR products were visualized on a 1% TBE-agarose gel. PCR product was cleaned up using Agencourt AMPure XP and submitted for cycle sequencing. Sequences have been submitted to GenBank XXXXXX.

#### Immunohistochemistry - opsin relative distribution and density

For the IHC assay, eyes were fixed overnight at 4°C in phospho-buffered saline (PBS) 4% paraformaldehyde (PFA), transferred into 1% PBS, and stored in 1% PFA at 4°C in 1% PBS until further processing. We supplemented the IHC sample by obtaining eyes from 34 additional bat species from the alcohol collections of the AMNH. As these specimens had been stored in 70% ethanol, once excised tissues were rehydrated in 1% PBS prior to dissection and then stored as above. Prior to processing, retinas were dissected from the eyeballs and were flattened by making three or four radial incisions from the outside of the retina inwards, with the deepest cut in the nasal pole. Immunodetection was carried out following standard procedures described in Ortin-Martinez *et al.* (2014). Retinas were permeated with two washes in PBS 0.5% Triton X-100 (Tx) and frozen for 15 minutes at –70°C in 100% methanol. Retinas were then thawed at room temperature, rinsed twice in PBS-0.5%Tx and incubated overnight at 4°C in the appropriate mixture of primary antibodies diluted in blocking buffer (PBS, 2% normal donkey serum, 2%Tx). The next day, retinas were washed four times in PBS-0.5Tx and incubated for 2 hours at room temperature in secondary antibodies diluted in PBS-2%Tx. Finally, retinas were thoroughly washed four times in PBS-0.5%Tx and, after a last rinse in PBS, mounted scleral side up on slides in anti-fading solution (Prolong Gold Antifade, Thermofisher). For each species, IHC was performed on at least three retinas from at least three field-collected individuals, and three retinas from two individuals for museum-sampled species (for details see Table S1). Given the consistency of the detection of these antibodies across all bats and other mammals (Müller et al., 2009; Müller et al., 2007; Ortín-Martínez et al., 2014), and the number of replicates and individuals examined, we interpreted no labeling to indicate a true loss of the respective cone type.

#### IHC - Antibodies and working dilutions

The following primary antibodies were used: goat anti-OPN1SW, 1:1000 (sc-14363, Santa-Cruz Biotechnologies, Heidelberg, Germany; detects S-opsin protein) and rabbit anti-opsin red/green, 1:750 (ab5405, Millipore Ibérica, Madrid, Spain; detects L-opsin protein). The following secondary antibodies were used at a 1:500 dilution: donkey antigoat Alexa Fluor 568 and donkey anti-rabbit Alexa Fluor 647 (Thermofisher).

#### IHC - Photoreceptor quantification

Flat-mounted retinas were photographed using a 20× objective on a confocal microscope (LSM710; Zeiss Microscopy). 564 and 633 lasers were used to excited Alexa 568 and Alexa 647 dyes, each labelling respectively S- and L-opsins. Each entire retina was completely imaged using 512×512 pixel tiles. For each retina, each tile was then Z-stacked and automatically counted using a 3D object counter plugin using Fiji (ImageJ). The accuracy of this automatic approach was verified by manually counting three biological replicates of five bat species, by two different people. For each retina quantified, the density was calculated for each tile and then averaged for each individual (total count was average over three individuals) and for each species (by averaging the average of the three individuals). The spatial distribution of L- and S-cone density was visualized for the following 14 species: *Mormoops blainvillei*, *Pteronotus quadridens*, *Macrotus waterhousii*, *Desmodus rotundus*, *Gardnerycteris crenulatum*, *Monophyllus redmani*, *Glossophaga soricina*, *Erophylla sezekorni*, *Brachyphylla nana pumila*, *Carollia sowelli*, *Sturnira lilium*, *Artibeus jamaicensis*, *Artibeus phaeotis*, *and Phyllops falcatus* (see Figure 3, Table S4).

#### Opsin gene evolution

We obtained protein-coding sequences for *OPN1LW* and *RHO* from the transcriptomes of all 38 species and included sequence data for *Eptesicus fuscus* (from GenBank), for a total of 39 species. For *OPN1SW*, we obtained sequence data for 28 species from the transcriptome data sets. We then supplemented the *OPN1SW* nucleotide sequences with those extracted from genome data using a combination of blastn and bl2seq on five noctilionoid bat genomes (*Artibeus jamaicensis*, *Desmodus rotundus* (Zepeda Mendoza et al., 2018), *Lionycteris spurrelli*, *Macrotus waterhousii*, *Mormoops blainvillei* and *Noctilio leporinus*) using the *Miniopterus natalensis OPN1SW* (XM_016213323.1) sequence as a query. The *L. spurrelli* and *M. waterhousii* genomes were sequenced by the Rossiter Lab, and the *A. jamaicensis*, *M. blainvillei* and *N. leporinus* genomes were made available by the Broad Institute. We also obtained the *OPN1SW* sequences from *E. fuscus* from GenBank. The extracted and aligned sequences are available from DRYAD XXXX.

Sequences for *OPN1LW* and *RHO* were aligned using MUSCLE v3.8.425 (Edgar, 2004) as translated amino acids to keep the sequences in frame. The software implementation of our model requires the alignment to be in frame without any stop codons, therefore, stop codons at the end of the reading frame were removed from the alignment. Due to stop codons and indels, nucleotide sequences for *OPN1SW* were aligned by eye, with columns containing disruptions to the reading frame being removed to keep the remaining sequences in frame. Hypervariable regions at the beginning or end of sequences that may be caused assembly errors were masked by ‘Ns’ in two cases (*Phyllops falcatus* and *Phyllonycteris poeyi*). Additionally, premature stop codons were masked with ‘Ns’ and columns containing insertions that shifted the translation frame were deleted to keep codons in frame.

We tested for whether there were differences in rates of molecular evolution in the three opsin genes by estimating the ratio of the rates of nonsynonymous to synonymous substitutions (*ω*) for different branch classes. We set up two frameworks: S-cone presence and diet. When looking at S-cone presence variation, in our 2-branch class test, we estimated differences for branches that lacked the S-cone protein (*ω*_S-cone.absent_), and those that had the S-cone protein present (*ω*_background_). We also designed a three branch-class test in which we estimated different rates for bats with Scones (*ω*_background_), bats that lack S-cones but have an intact reading frame for the *OPN1SW* transcripts (*ω*_OPN1SW.intact_), and bats that lack S-cones and *OPN1SW* is a pseudogene (*ω*_OPN1SW.pseudo_). If bats that lack the S-cone experience relaxed selection, we expect higher rates of *ω* in bats without the S-cone in the *OPN1SW*gene, but no differences among groups in the other two opsins. The sequences from lineages for which no S-cone data was available were removed from this analysis (*n* = 8). Finally, we tested if there were differences in rates in frugivorous lineages (*ω*_frugivore_) and all other bats (*ω*_background_). Figure S2 depicts branches labeled with respective branch classes. Frugivory data was available for all lineages, and thus all available sequences were used in this analysis.

These analyses were performed using the branch model implemented in the codeml routine of PAML 4.8a (Yang, 2007). Differences among branches were compared against estimates for a single *ω* for all branches (*ω*_background_). We used a likelihood ratio test to compare the best-fit model for each opsin gene. The analysis used the species topology that merged a recently published phylogeny of all bats (Shi & Rabosky, 2015) with that of a recently published noctilionoid tree (Rojas et al., 2016). The tree was trimmed using the geiger v. 2.0.6 package in R (Harmon et al., 2008).

#### Ecological correlates of cone presence and density

A hierarchical Bayesian approach was used to quantify the influence of ecology on the presence of S-cones while accounting for the phylogenetic correlation between observations from different species. A hierarchical approach is often called a mixed model in the literature, with cluster-specific effects called “random”, and sample-wide effects called “fixed”. As different fields apply random and fixed to different levels of the hierarchy, here we adopt the language of cluster-specific and sample-wide effects (Gelman, 2005). The effect of species was quantified by including species as a cluster-specific, or random effect in the R package MCMCglmm (Hadfield, 2010). Additionally, to address variation among different estimates of phylogeny, we used the R package mulTree (Guillerme & Healy, 2014) to run the Bayesian models across a sample of trees obtained from the posterior of phylogenies.

The first set of models explained presence or absence of S-cones as a series of ecological dummy variables coded as prevalence or non-prevalence of plant materials, fruit, or other (insects or small vertebrates) items in diet. In the sample-wide or fixed portion of these models, observations *y* for each species from 1 to *i* for each diet category 1 to *j* correspond to a single-trial binomial response of the probability of observing S-cones given by *pr_i_* such that:

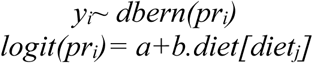

Models for each of the dietary categorizations were then compared using the Deviance Information Criterion or DIC. For Bayesian models, a lower DIC can indicate a better fit of the model to the data (Gelman, 2004). The predictor variable identified by the best-fit model for presence/absence of S-cones was then used in subsequent analyses of cone density.

Analyses of the sample-wide or fixed portion of cone density modeled the natural logarithm of the cone density *y* for each observation *i* as a function of dummy predictor variables defined by an diet group *j*, or S-cone group *k*. *ln(yi*) was modeled as a random, normally distributed variable with mean mu and variance, as below:

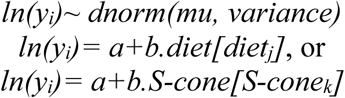

Unlike the presence/absence analyses, these response variables were normally distributed, with the sample-wide portion of the model accounting for the effect of diet or S-cone presence on L-cone density. The cluster-specific or random effect accounted for both the relationships between species and the clustering of observations when more than one measurement was taken for each species.

To estimate the covariation arising from phylogeny for all species analyzed, we used the phylogeny of Shi and Rabosky (2015) for non-noctilionoids and as a base tree. For New World noctilionoids, a posterior sample of 100 trees from the New World noctilionoid phylogenies of Rojas *et al.* (2016) was grafted onto the base tree and used as input in mulTree analyses. Each regression corresponding to one of the 100 phylogenies ran for 20M generations, sampling every 1000 generations, with a burn-in of 100,000. Each regression ran 2 separate chains, assessing post-burnin convergence by comparing posteriors and reaching estimated sampling sizes (ESS) of at least 200 for all model parameters.

MCMCglmm uses inverse Wishart distributions for priors on sample-wide variance and cluster-specific or phylogenetic variance. For uncorrelated predictor variables, these functions collapse into inverse gamma distributions for residuals. The choice of the residual prior is particularly important for phylogenetic logistic regressions (Ives & Garland, 2014), for which this variance is not identified in the likelihood (Hadfield, 2010). Hence, to properly estimate the posterior, we fixed the residual variance at 1. As the phylogenetic structure of the data corresponds to a matrix structure of correlations between observations, it becomes necessary to expand the parameters by specifying both the mean and covariance matrix on the prior, in addition to the shape and scale parameters given by nu/2. For the logistic regressions, the prior on the phylogenetic structure was given by nu = 1000 and *V* = 1 (generating a very long-tailed distribution), with prior mean of 0 and covariance of 1. For Gaussian regressions, the prior on the residuals was given by nu = 1 and *V* = 1 (Hadfield, 2016).

The results from all 100 posterior trees were combined to generate parameter estimates and 95% high-probability density from the joint posterior probability distribution. Predictors were found to influence the response if the high-probability density for their coefficient did not overlap with 0. We also calculated the explained variance or *R*^2^ based on the method proposed by Nakagawa and Schielzeth (2013). Two types of *R*^2^ were calculated for hierarchical Bayesian models, based only on the sample-wide factors or marginal *R*^2^, and based on both sample-wide and cluster specific factors, or conditional *R*^2^.

### Supplementary Results

#### RNA quality and sequencing statistics

Integrity of extracted RNA varied across samples (RIN 4.1 to 10), the majority of samples obtained RINs greater than the recommended value of 8. Following sequencing, the number of raw reads ranged from 9,129,366 to 47,639,831 per sample. Cleaned reads ranged from 8,659,526 to 45,280,391 per sample, which resulted in assemblies of 67,459 to 131,925 across species.

For three species (*Glossophaga soricina*, *Vampyrodes caraccioli* and *Artibeus bogotensis*), the *OPN1SW* transcript was recovered following chimera removal as the transcript was initially joined to that of *CALU.* The *OPN1SW*transcript recovered for *Carollia brevicauda* contained an internal stop codon, however, this was not shared by *Carollia perspicillata.* Mapping of the *C. brevicauda* short reads against *C. perspicillata* using the script align_and_estimate_abundance.pl available from Trinity utils, and RSEM and bowtie, suggested low-coverage and mapped reads did not show this mutation, thus this mutation might not be genuine. Manual blasting of the *Phyllonycteris poeyi* and *Erophylla bombifrons* assemblies for *OPN1SW* recovered several fragmentary sequences that could represent this gene. However, sections of the sequences were highly divergent and were found to represent intronic regions by BLAST implying there could be retained introns in the mRNA.

#### Immunohistochemistry

We used IHC to assess the presence or absence of OPN1SW and OPN1LW proteins in whole, flat-mounted retinas of adult bats (Figures 2 and S1). We detected OPN1LW proteins in the adult eyes of all sampled bat species (*n* = 56 species total), whereas we detected the OPN1SW protein in the eyes of only 32 bat species. The following species lack the OPN1SW protein: *Molossus molossus, Eptesicus fuscus, Chilonatalus micropus, Pteronotus davyi, P. parnellii, Mormoops blainvillei, Macrotus waterhousii, Diaemus youngi, Desmodus rotundus*, *Trachops cirrhosus*, *Tonatia saurophila*, *Gardnerycteris crenulatum*, *Monophyllus redmani*, *Erophylla sezekorni*, *E. bombifrons*, *Brachyphylla pumila*, *Lonchophylla robusta*, and *Carollia brevicauda.* Specimens assayed for *Phyllostomus hastatus*, *Sturnira tildae*, *S. ludovici*, and *Platyrrhinus dorsalis*, were obtained from museum specimens. While we detected the OPN1LW protein in these samples, they were characterized by a low signal-to-background ratio in the OPN1SW protein labeling, which prevented us from determining OPN1SW presence or absence.

In most species examined either both the *OPN1SW* transcript and protein were detected (*n* = 32), or neither were detected (*n* = 5). For example, all members of the Stenodermatinae clade of fruit-eating bats examined were found to have both the *OPN1SW* transcript and protein. Outside of this clade, we also detected the OPN1SW protein, and in most cases confirmed transcript presence with RNA-Seq, in species distributed widely throughout the phylogeny. Including within Emballonuridae (*Saccopteryx bilineata* and *S. leptura*), Noctilionidae (*Noctilio leporinus*), Mormoopidae (*Pteronotus quadridens*), and other Phyllostomidae (*Phyllostomus elongatus*, *Anoura geoffroyi*, *Glossophaga soricina*, *Carollia perspicillata*, *Rhinophylla fischerae*, and R. *pumilio*).

We detected the *OPN1SW* transcript, but no protein, in fewer species. These species are also widely distributed throughout the phylogeny and include the following: *Molossus molossus*, *Pteronotus parnellii*, *Tonatia saurophila*, *Gardnerycteris crenulatum*, *Monophyllus redmani*, *Erophylla bombifrons*, and *Carollia brevicauda.*

#### Opsin gene evolution

The alignments used for input for the PAML analyses resulted in 349 codons for *OPN1SW*, 364 for *OPN1LW*, and 348 for *RHO.* For the analysis testing the difference in *ω* rates between species that do and do not express the S-cone protein, there was no difference in *ω* estimates between branch classes for the *RHO* (*χ*^2^_(1)_ = 0.12, *P* = 0.73; *χ*^2^_(2)_ = 0.41, *P* = 0.81). However, there was a difference favoring the three-branch class model for *OPN1SW* (*ω*_background_ = 0.11; *ω*_OPN1SW.intact_ = 0.07; *ω*_OPN1SW.pseudo_ = 0.79; *χ*^2^_(2)_ = 129.9, *P* < 1.0e-5) and *OPN1LW* (*ω*_background_ = 0.08; *ω*_OPN1SW.intact_ = 0.08; *ω*_OPN1SW.pseudo_= 0.17; *χ*^2^_(2)_ = 6.9, *P* = 0.03) gene (Table S2). For the analysis testing for difference in *ω* rates between frugivorous species and background branches, there was no difference in *ω* estimates between branch classes for the *OPN1SW* (*χ*^2^_(1)_ = 0.4, *P* = 0.53) and *OPN1LW* (*χ*^2^_(1)_ = 0.49, *P* = 0.48) genes (Table S3). However, there was a difference in *ω* for the *RHO* gene (Table S3), in frugivorous lineages showed significantly lower rates than background branches (*ω*_background_ = 0.04; *ω*_fugivory_= 0.02; *χ*^2^_(1)_ = 11.0, *P* = 9.1e-4).

#### Ecological correlates of cone presence and density

Tables S5-S7 and Figure S4 summarize the results from Bayesian regressions. The frequency distributions of the DIC for three sets of phylogenetic regressions of S-cone presence against particular diet prevalence reveal the prevalence of fruit results in the least DIC of any model, providing the best fit to these data (Fig. S4). The posterior estimates of parameters for this model reveal the prevalence of fruit in diet increases the odds of having the S-cone by a factor of 37.7 (given by *e*^^^coefficient of fruit prevalence, or from odd of ∼0.34 to ∼12.8, Table S5). Analyses of the density of long-wave cones revealed no statistically meaningful effect of fruit prevalence on this density (Table S6). Instead, analyses of the density of long-wave cones as a function of the presence of S-cones revealed having the S-cones increases the density of L-cones by 0.43 in the natural logarithm scale (or a factor of ∼1.54 in the linear scale) compared to species without S-cones (or from a baseline of ∼3944 to ∼6063, Table S7).

## Supplementary Tables and Legends

**Table S1.**
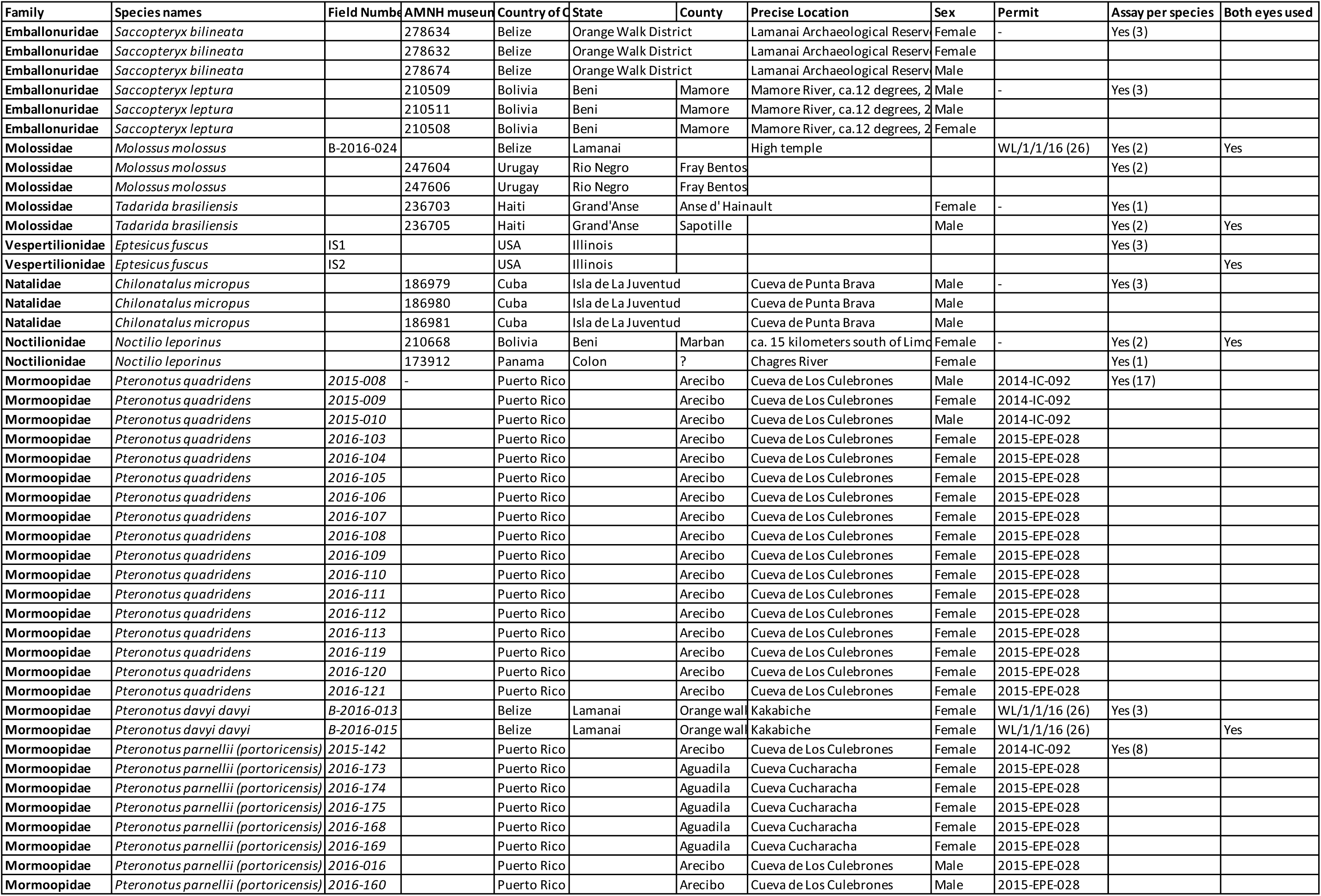
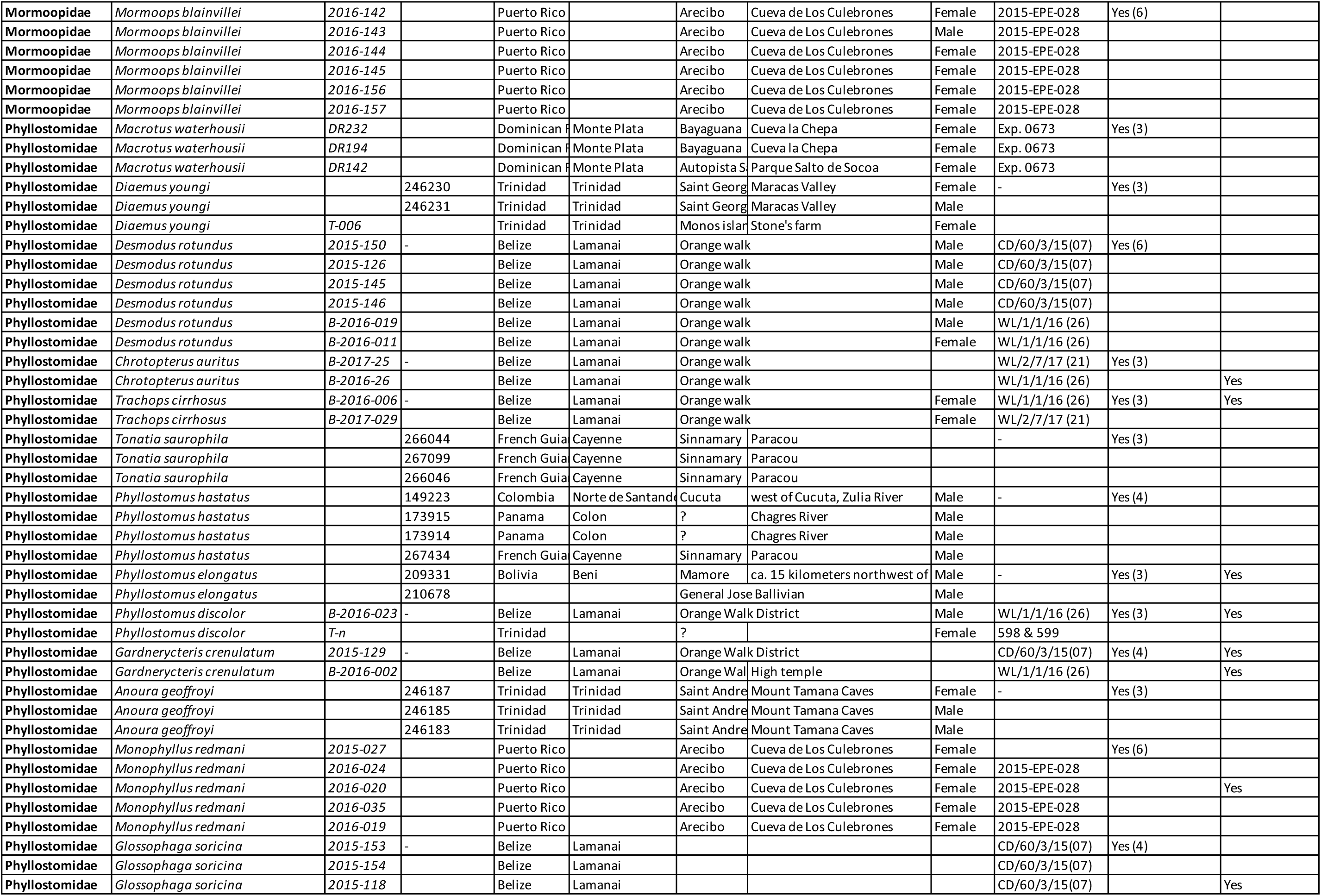
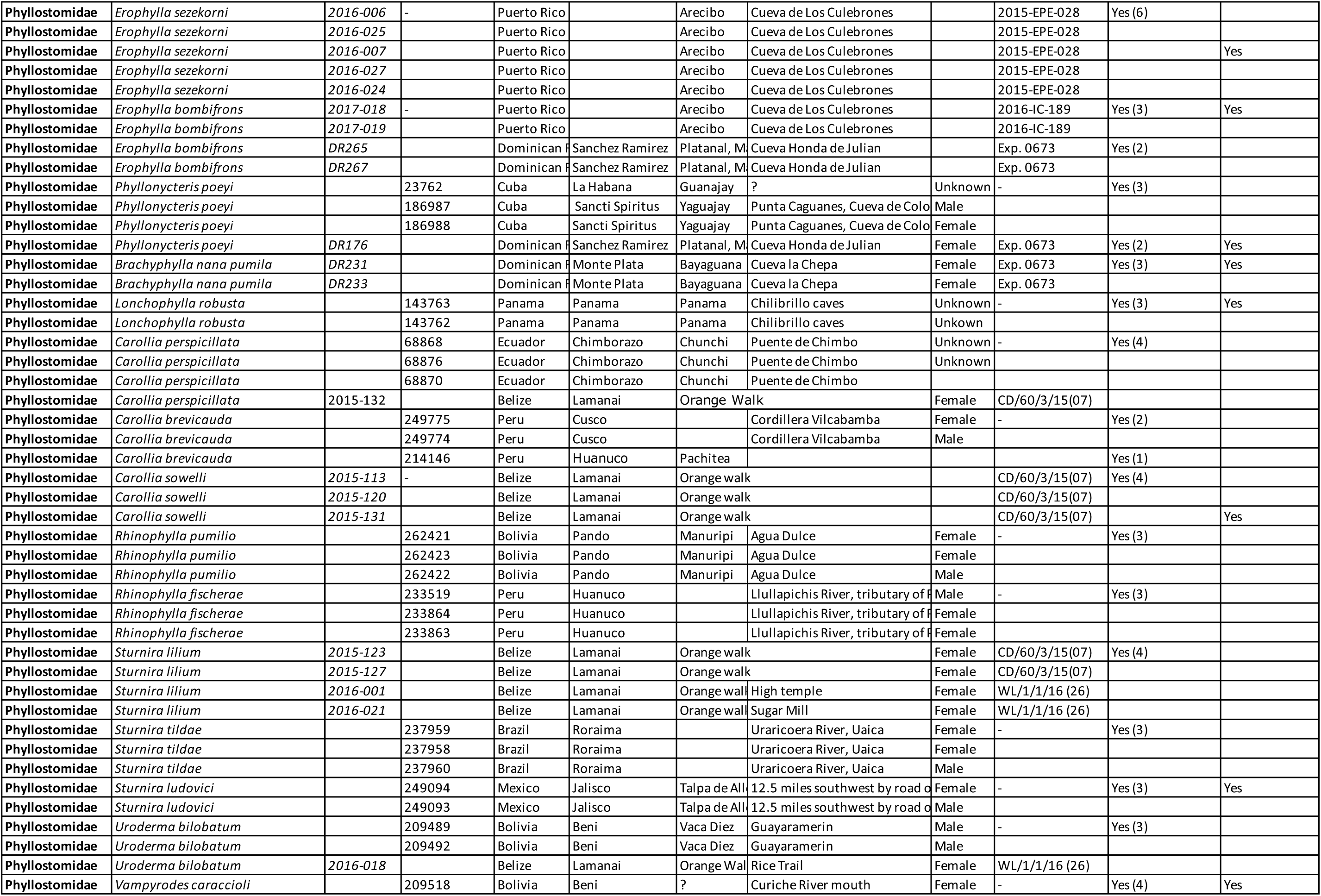
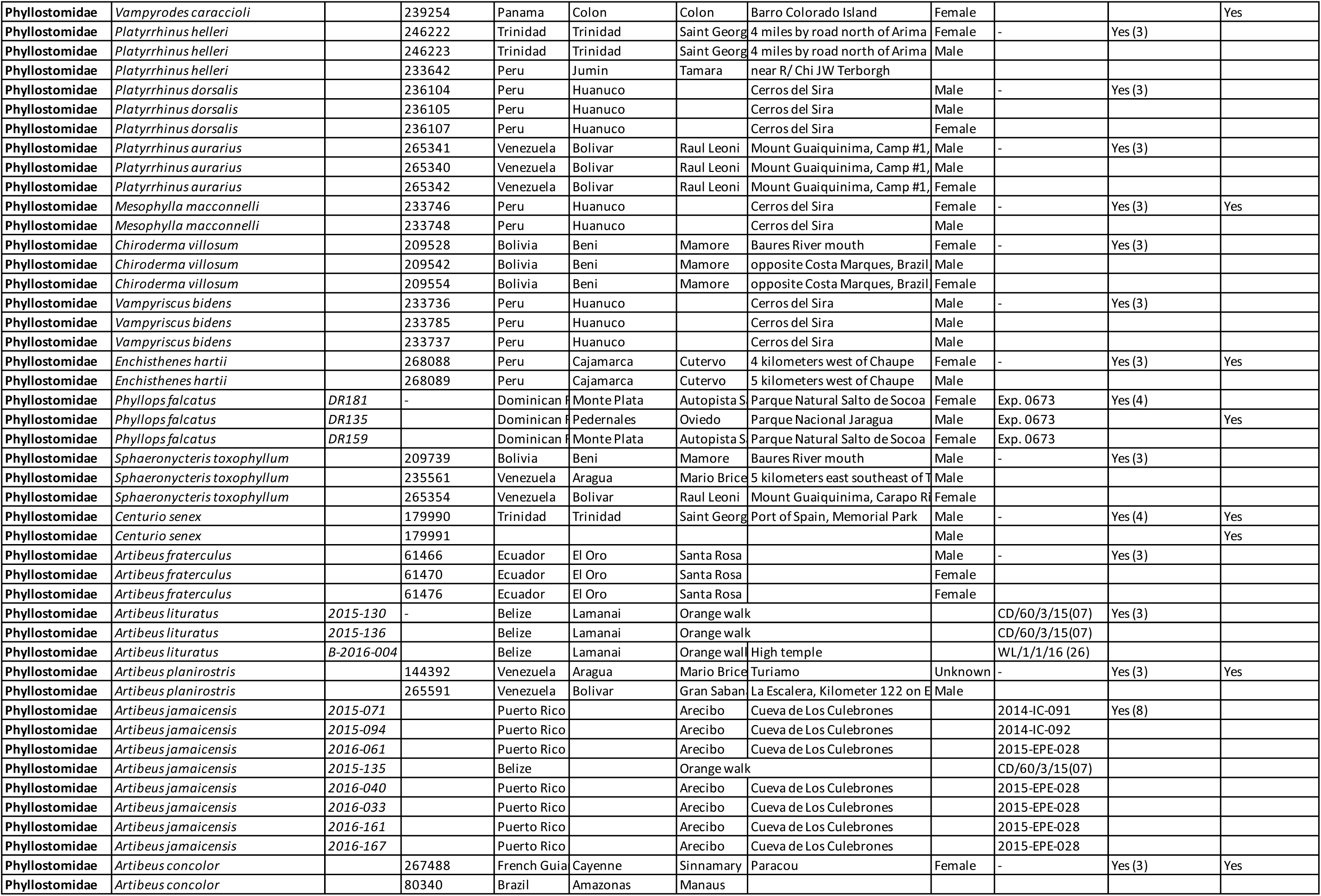
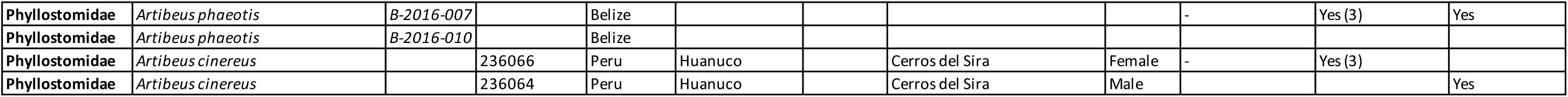
– as separate Excel file. Specimen and sampling information for the tissues used by this study.

**Table S2.** Results of molecular evolution branch analyses for each of the three opsin genes tested for differences in rates of nonsynonymous to synonymous substitutions (*ω*) for lineages that lack the S-cone and lineages that have retained the S-cone. Grey boxes indicate the preferred model inferred from a likelihood ratio test. *lnL*: log-likelihood; *np*: number of parameters; *TL*: tree length; *k*: kappa (transition/transversion ratios); LR: likelihood ratio; *p*: *p*-value of likelihood ratio of alternative relative to null for each test

| Gene | Model | lnL | np | TL | k | LR | p | $\omega$ |
| --- | --- | --- | --- | --- | --- | --- | --- | --- |
| **OPN1SW** | *null* | -5176 | 57 | 2.41 | 4.7 | --- | --- |  |
| | | | | | | | $\omega_{\text{background}}$ | 0.19 |
|  | *2 branch* | -5156 | 58 | 2.41 | 4.7 | 47.8 | < 1.0e-5 |  |
| | | | | | | | $\omega_{\text{background}}$ | 0.11 |
| | | | | | | | $\omega_{\text{S-cone.absent}}$ | 0.35 |
|  | *3 branch* | -5111 | 59 | 2.41 | 4.7 | 129.9 | < 1.0e-5 |  |
| | | | | | | | $\omega_{\text{background}}$ | 0.11 |
| | | | | | | | $\omega_{\text{OPN1SW.intact}}$ | 0.07 |
| | | | | | | | $\omega_{\text{OPN1SW.pseudo}}$ | 0.79 |
| **OPN1LW** | *null* | -4124 | 61 | 1.45 | 6.7 | --- | --- |  |
| | | | | | | | $\omega_{\text{background}}$ | 0.10 |
|  | *2 branch* | -4123 | 62 | 1.45 | 6.7 | 2.77 | 0.10 |  |
| | | | | | | | $\omega_{\text{background}}$ | 0.08 |
| | | | | | | | $\omega_{\text{S-cone.absent}}$ | 0.12 |
|  | *3 branch* | -4121 | 63 | 1.45 | 6.7 | 6.9 | 0.03 |  |
| | | | | | | | $\omega_{\text{background}}$ | 0.08 |
| | | | | | | | $\omega_{\text{OPN1SW.intact}}$ | 0.08 |
| | | | | | | | $\omega_{\text{OPN1SW.pseudo}}$ | 0.17 |
| **RHO** | *null* | -4101 | 61 | 1.8 | 7.6 | --- | --- |  |
| | | | | | | | $\omega_{\text{background}}$ | 0.03 |
|  | *2 branch* | -4101 | 62 | 1.8 | 7.6 | 0.35 | 0.55 |  |
| | | | | | | | $\omega_{\text{background}}$ | 0.03 |
| | | | | | | | $\omega_{\text{S-cone.absent}}$ | 0.03 |
|  | *3 branch* | -4101 | 63 | 1.8 | 7.6 | 0.41 | 0.81 |  |
| | | | | | | | $\omega_{\text{background}}$ | 0.03 |
| | | | | | | | $\omega_{\text{OPN1SW.intact}}$ | 0.03 |
| | | | | | | | $\omega_{\text{OPN1SW.pseudo}}$ | 0.03 |

**Table S3.**
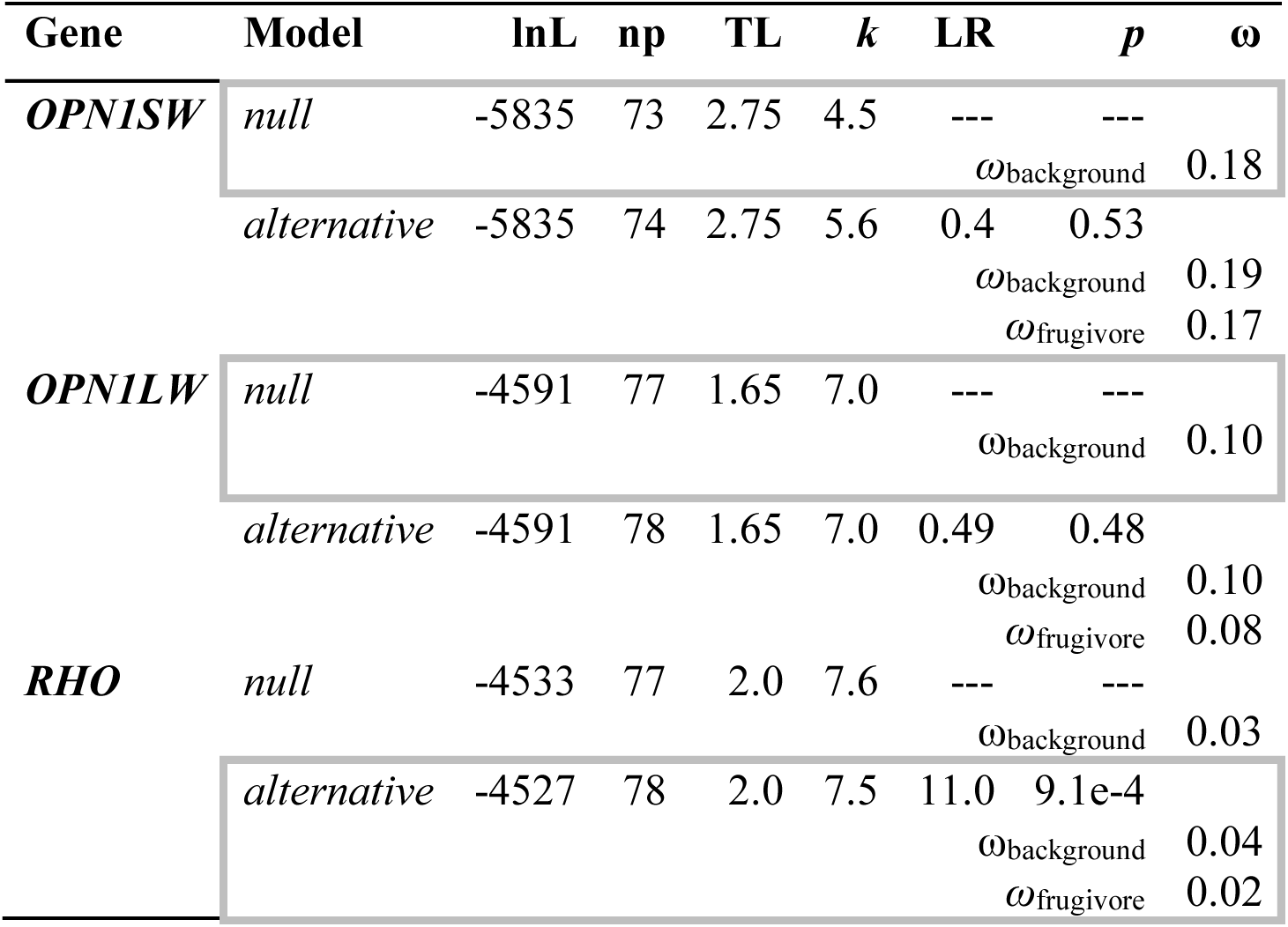
Results of molecular evolution branch analyses for each opsin gene that test for differences in rates of nonsynonymous to synonymous substitutions (*ω*) for frugivorous and non-frugivorous lineages. Grey boxes indicate the preferred model inferred from a likelihood ratio test. *lnL:* log-likelihood; *np:* number of parameters; *TL:* tree length; *k*: kappa (transition/transversion ratios); LR: likelihood ratio; *p*: *p*-value of likelihood ratio of alternative relative to null for each test

**Table S4.** Cone densities in 14 species representative of different diet types. Mean cones densities across the retinal tiles quantified in individual flat mounted retinas. Densities were estimated after averaging the count for each tile.

| **Species** | **L-opsin** | **S-opsin** | **Field number** |
| --- | --- | --- | --- |
| *Artibeus jamaicensis* | 5534 | 2688 | 2015-071 |
| *Artibeus jamaicensis* | 5587 | 2253 | 2016-040 |
| *Artibeus phaeotis* | 5557 | 3642 | 2016-010 |
| *Artibeus phaeotis* | 6539 | 3841 | 2016-007 |
| *Carollia sowelli* | 4417 | 1094 | 2015-131 |
| *Carollia sowelli* | 4769 | 1162 | 2015-113 |
| *Carollia sowelli* | 4706 | 1178 | 2015-113 |
| *Carollia sowelli* | 4672 | 1194 | 2015-120 |
| *Sturnira lilium* | 4918 | 952 | 2015-123 |
| *Sturnira lilium* | 5109 | 1015 | 2015-127 |
| *Monophyllus redmani* | 2557 | 0 | 2015-027 |
| *Monophyllus redmani* | 2659 | 0 | 2015-020 |
| *Monophyllus redmani* | 2691 | 0 | 2015-020 |
| *Erophylla sezekorni* | 3681 | 0 | 2016-025 |
| *Erophylla sezekorni* | 3974 | 0 | 2016-007 |
| *Erophylla sezekorni* | 3499 | 0 | 2016-007 |
| *Erophylla sezekorni* | 3674 | 0 | 2016-027 |
| *Glossophaga soricina* | 5583 | 328 | 2015-153 |
| *Glossophaga soricina* | 5020 | 350 | 2015-118 |
| *Brachyphylla nana pumila* | 5471 | 0 | DR231 |
| *Brachyphylla nana pumila* | 5481 | 0 | DR231 |
| *Desmodus rotundus* | 3406 | 0 | 2015-150 |
| *Desmodus rotundus* | 3986 | 0 | 2015-126 |
| *Desmodus rotundus* | 3980 | 0 | 2015-145 |
| *Pteronotus quadridens* | 7227 | 4263 | 2016-109 |
| *Pteronotus quadridens* | 8100 | 5022 | 2016-108 |
| *Pteronotus quadridens* | 8109 | 6652 | 2016-119 |
| *Pteronotus quadridens* | 6487 | 7053 | 2016-121 |
| *Mormoops blainvillei* | 4089 | 0 | 2016-142 |
| *Mormoops blainvillei* | 3869 | 0 | 2016-142 |
| *Macrotus waterhousii* | 6692 | 0 | DR194 |
| *Gardnerycteris crenulatum* | 2080 | 0 | 2015-129 |
| *Phyllops falcatus* | 5944 | 2716 | DR135 |

**Table S5.** Summary of Bayesian logistic regression models of the presence of SW-cones as a function of whether or not fruit is prevalent in the diet. HPD, high-probability density.

| **Parameter** | **Median** | **2.5% HPD** | **97.5% HPD** |
| --- | --- | --- | --- |
| Intercept | -1.09 | -3.18 | 0.51 |
| Fruit prevalent | 3.63 | 1.82 | 6.72 |
| Phylogenetic variance | 1.90 | 0.02 | 8.58 |
| Conditional $R^2$ | 0.58 | 0.18 | 0.83 |
| Marginal $R^2$ | 0.88 | 0.64 | 0.96 |

**Table S6.**
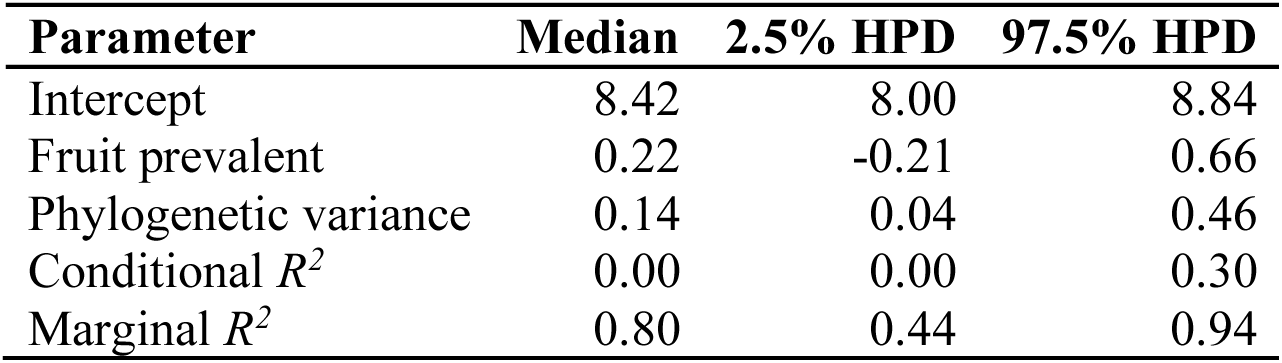
Summary of Bayesian regression models of the ln-transformed density of long-wave cones (plus one) as a function of whether or not fruit is prevalent in the diet. HPD, high-probability density.

**Table S7.** Summary of Bayesian regression models of the ln-transformed density of long-wave cones (plus one) as a function of whether or not they have S-cones. HPD, high-probability density.

| **Parameter** | **Median** | **2.5% HPD** | **97.5% HPD** |
| --- | --- | --- | --- |
| Intercept | 8.28 | 7.95 | 8.63 |
| S-cones present | 0.43 | 0.09 | 0.78 |
| Phylogenetic variance | 0.07 | 0.00 | 0.29 |
| Conditional $R^2$ | 0.24 | 0.00 | 0.51 |
| Marginal $R^2$ | 0.74 | 0.32 | 0.91 |

## Legends

**Supplementary Figure 1:**
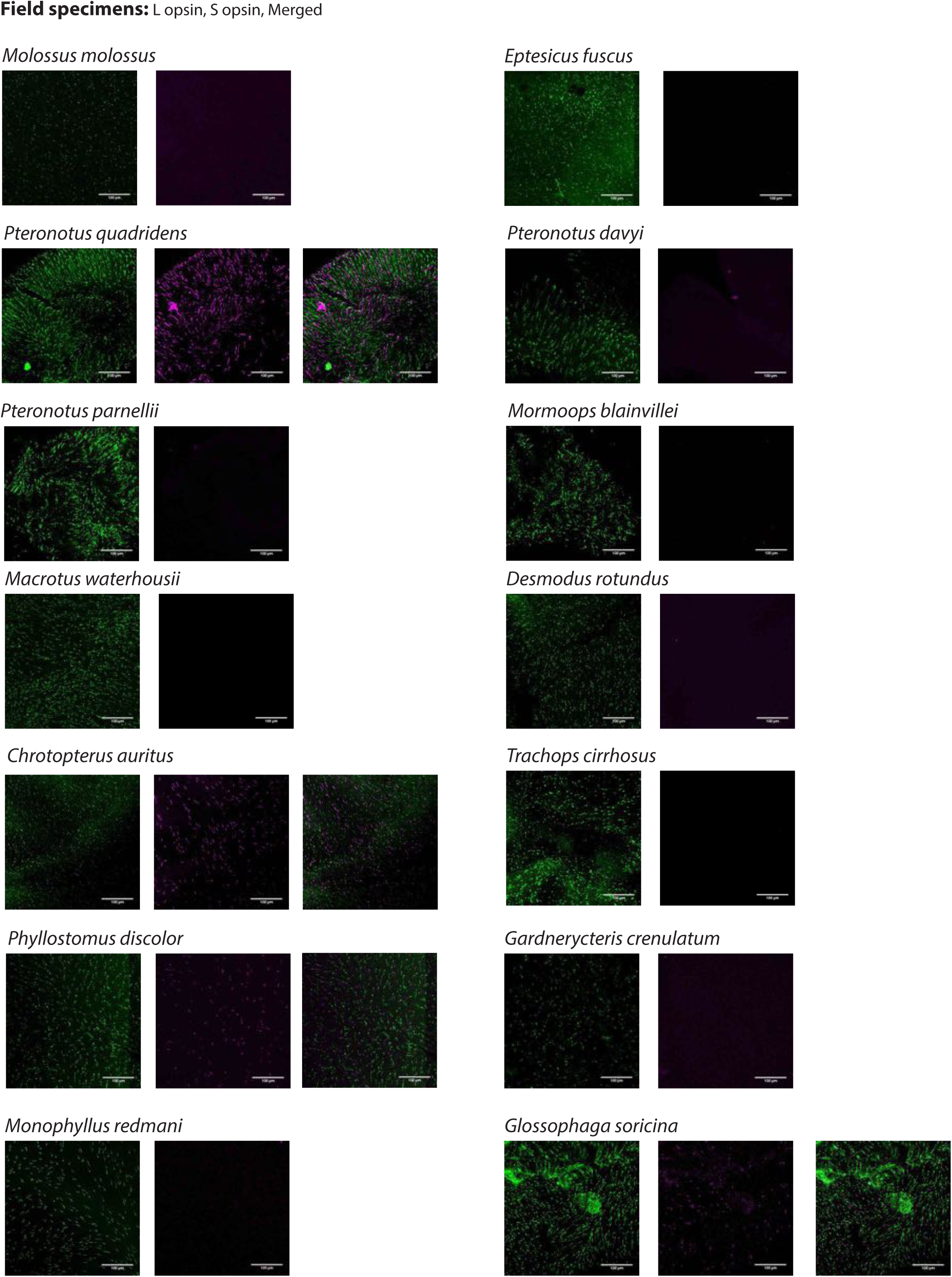
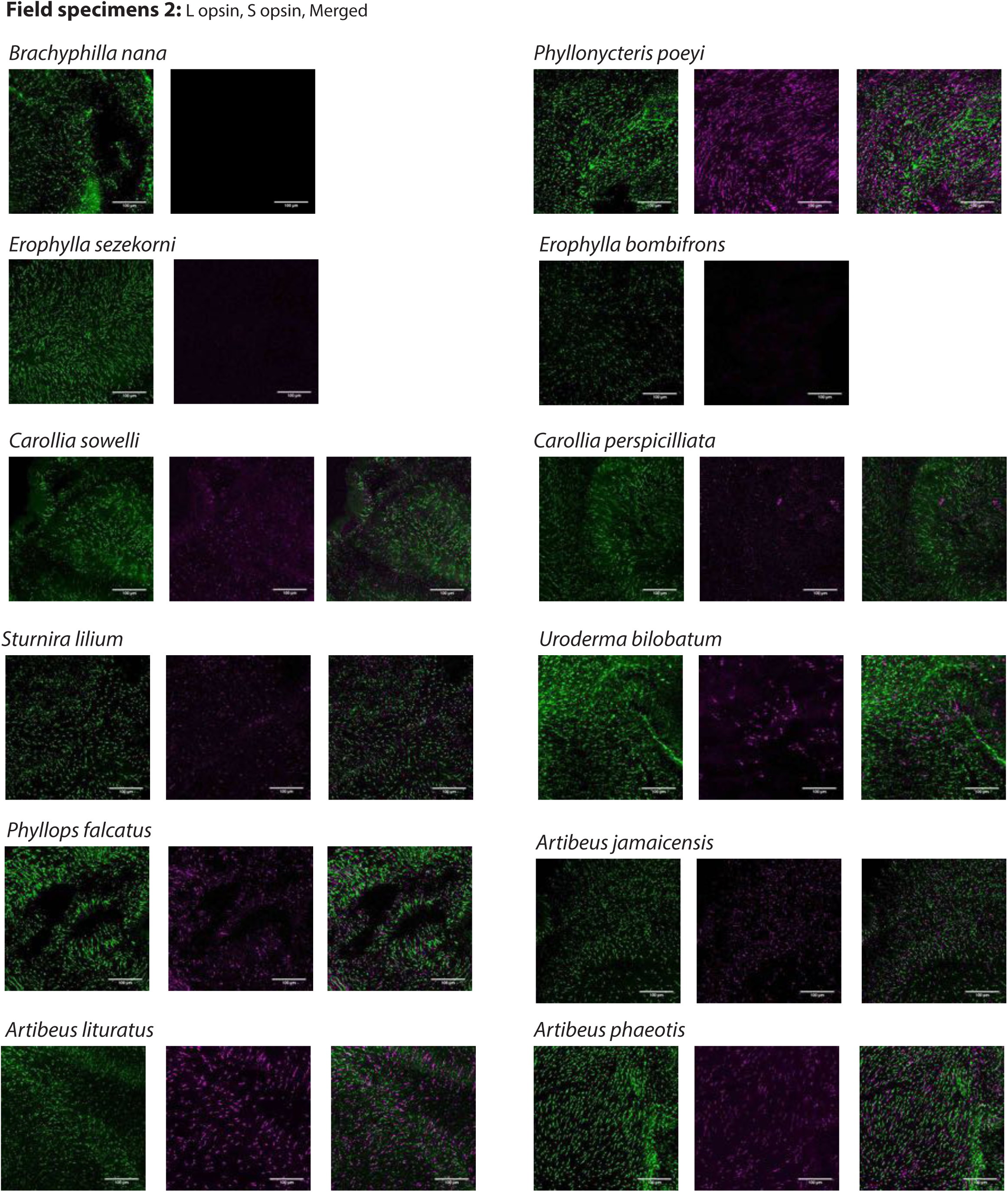
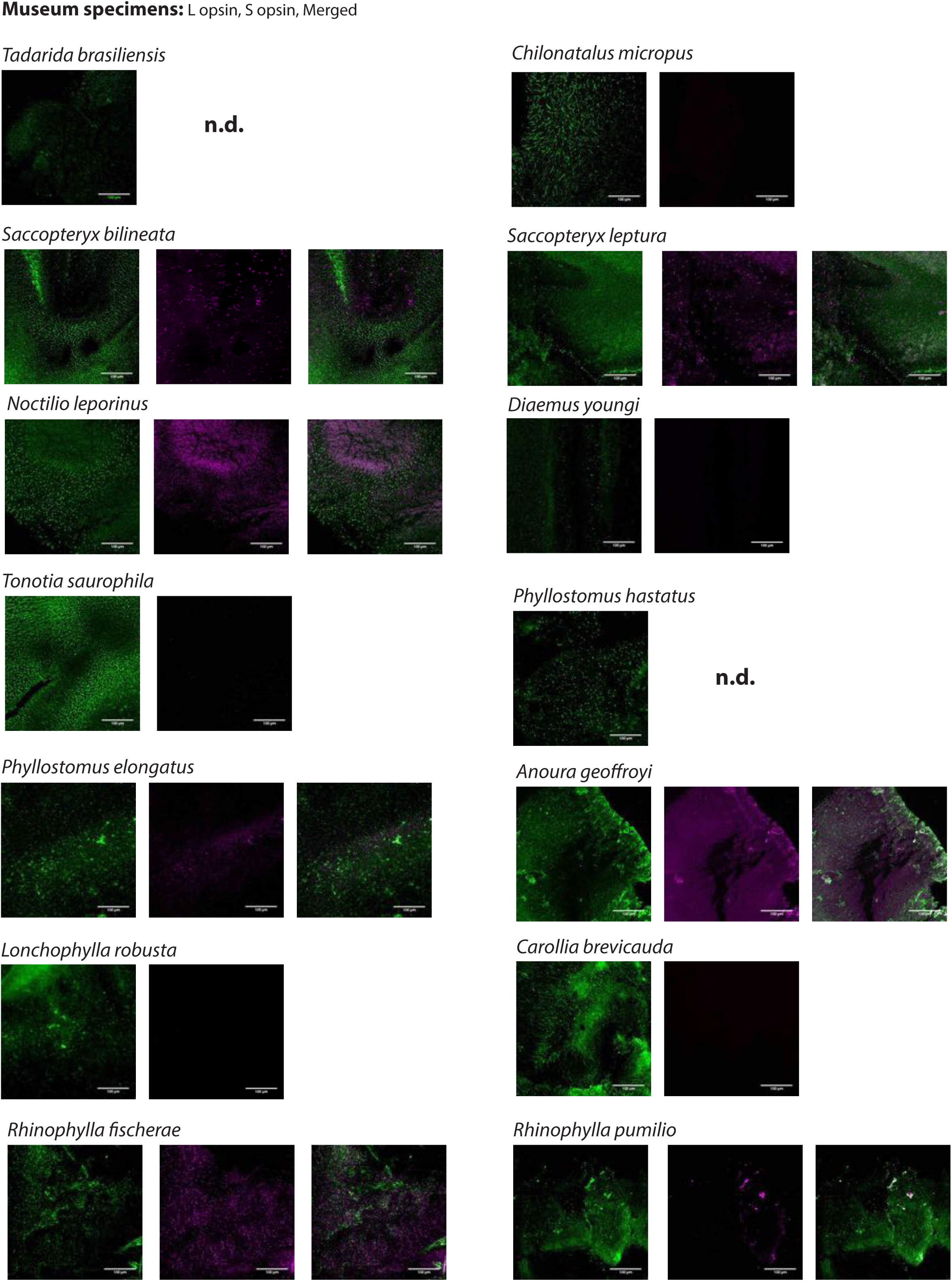
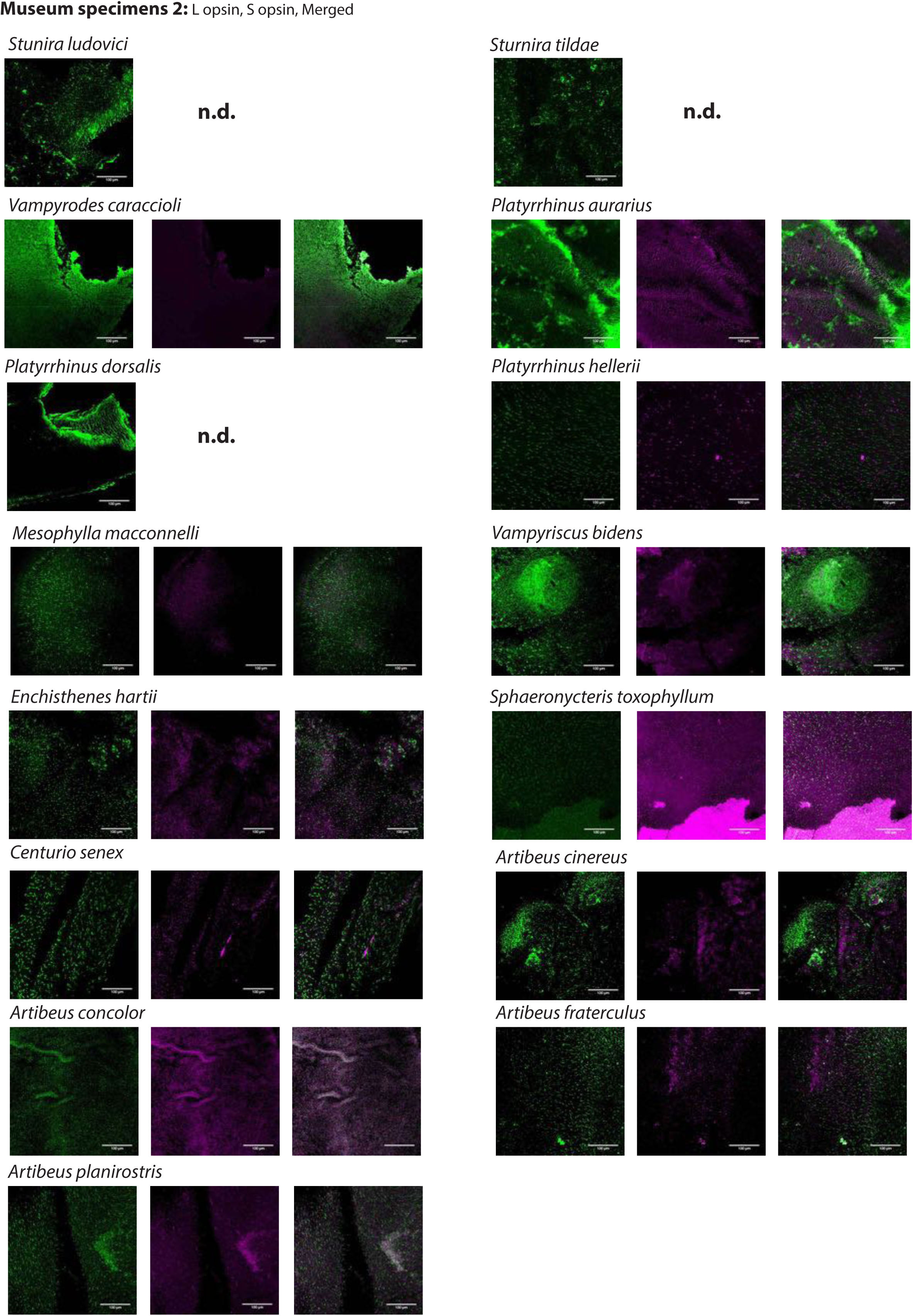
L- and S-opsin protein expression in L- and S-cones. L- and S-opsin presence was assayed by IHC in 55 species with antibodies recognizing each of L- and S-opsin. For each species, L- and S-cone labeling are shown. For species with both L- and S-cones, the merged column is the result of the merged images of L- and S-cones from the same individual. No evidence of dual cones was found. Scale bar: 100 μm.

**Supplementary Figure 2:**
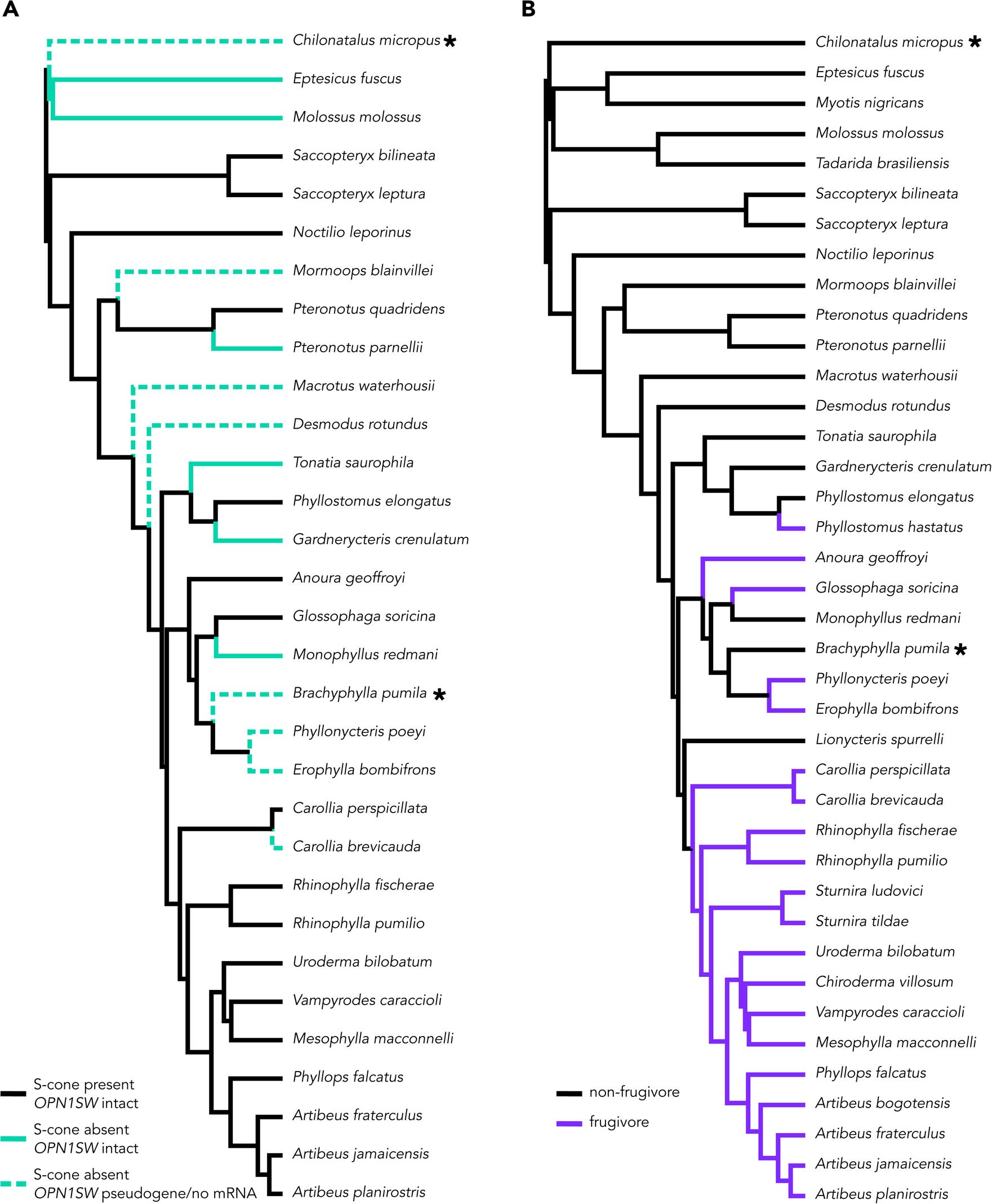
Branch class coding for molecular evolution analyses in PAML. Tests for S-cone variation and diet were run for all three genes. The ^∗^ indicates lineages absent from the *OPNSW1* alignment. (A) Branch coding for species with variations S-cone presence. In the two-branch class test, black branches (S-cone present) were compared against teal branches (S-cone absent). In the three-branch class test, teal branches were subdivided between lineages in which *OPN1SW* is intact (solid teal branch) and lineages in which *OPN1SW* is a pseudogene or the mRNA was absent (dashed teal branch). (B) Branch coding for frugivorous (purple) v. non-frugivorous lineages and background branches (black).

**Supplementary Figure 3:**
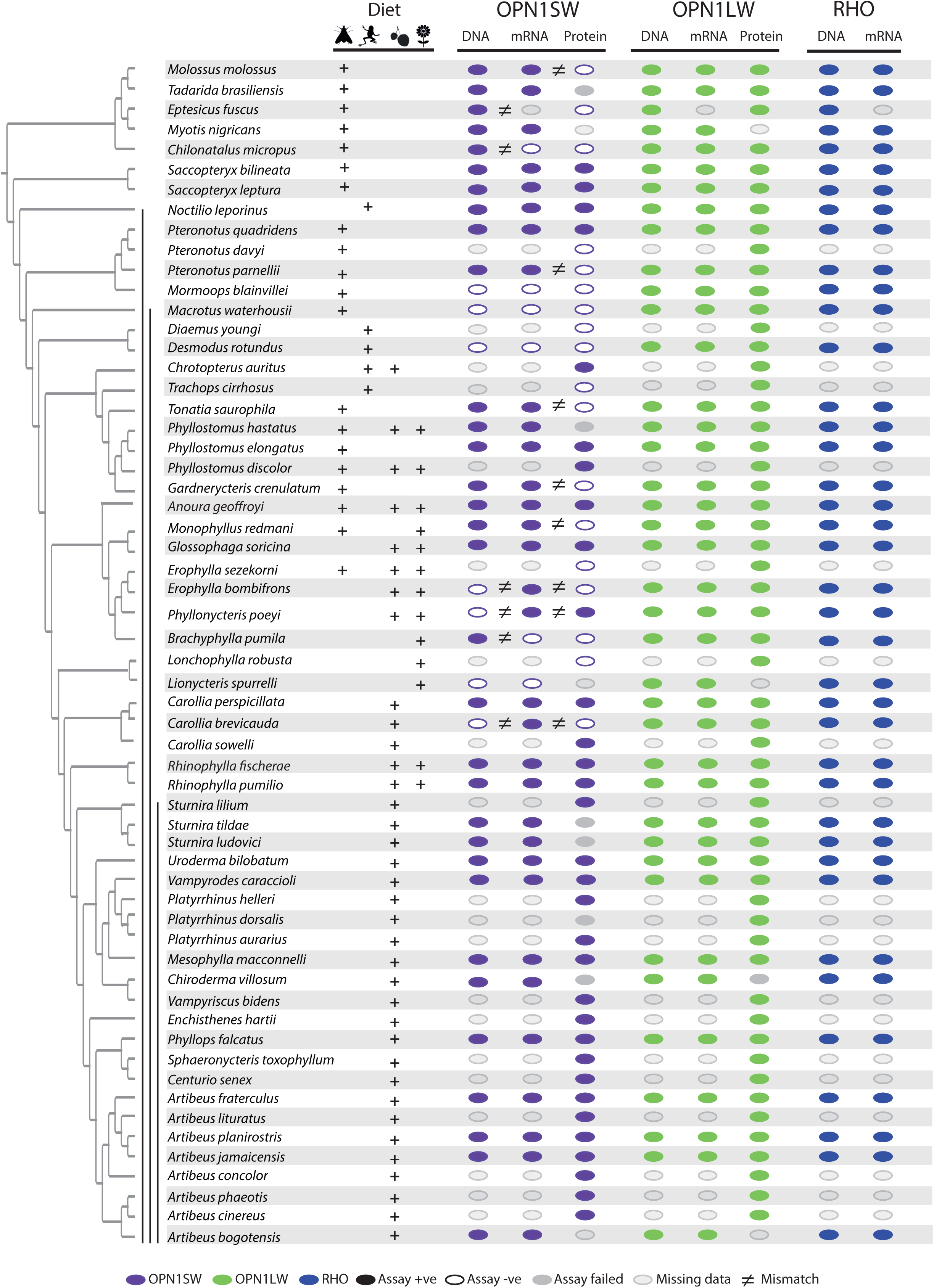
Distribution of an intact open reading frame (ORF), mRNA transcript and protein for the OPN1SW and OPN1LW photopigments in ecologically diverse noctilionoid bats. Diet composition follows Rojas et al. (2018), dietary types are indicated with the following symbols: invertebrates – moth, vertebrates – frog, fruit – fruit and nectar/pollen – flower. The species phylogeny follows (Rojas et al., 2016; Shi & Rabosky, 2015). Vertical black bars, from left to right, indicate: (1) Noctilionoidea, (2) Phyllostomidae, (3) and Stenodermatinae, respectively. RNA-Seq data was generated to both infer the presence of an intact ORF (in combination with genomic and PCR sequence data) and determine the presence of an expressed mRNA transcript. The presence of an intact ORF and mRNA transcript for RHO was verified across all transcriptomes. The presence/absence of a protein product for S- and L-opsins was assayed by IHC on flat mounted retinas. The presence of an intact ORF, mRNA and protein are indicated by a filled color marker, and its absence by an empty marker. Missing data (i.e. species for which we were unable to obtain tissue) are indicated with a grey marker with grey outline. Finally, a grey marker with no outline indicates the failure of protein assay for some species represented by museum specimens. Mismatches between intact ORFs and transcripts, or between transcripts and protein data are indicated by an inequality symbol. Note: OPN1SW protein assays for *Pteronotus quadridens* revealed polymorphisms within the sample, and we recorded positive OPN1SW assays in some *Phyllonycteris poeyi* individuals despite an apparent disrupted ORF.

**Supplementary Figure 4:**
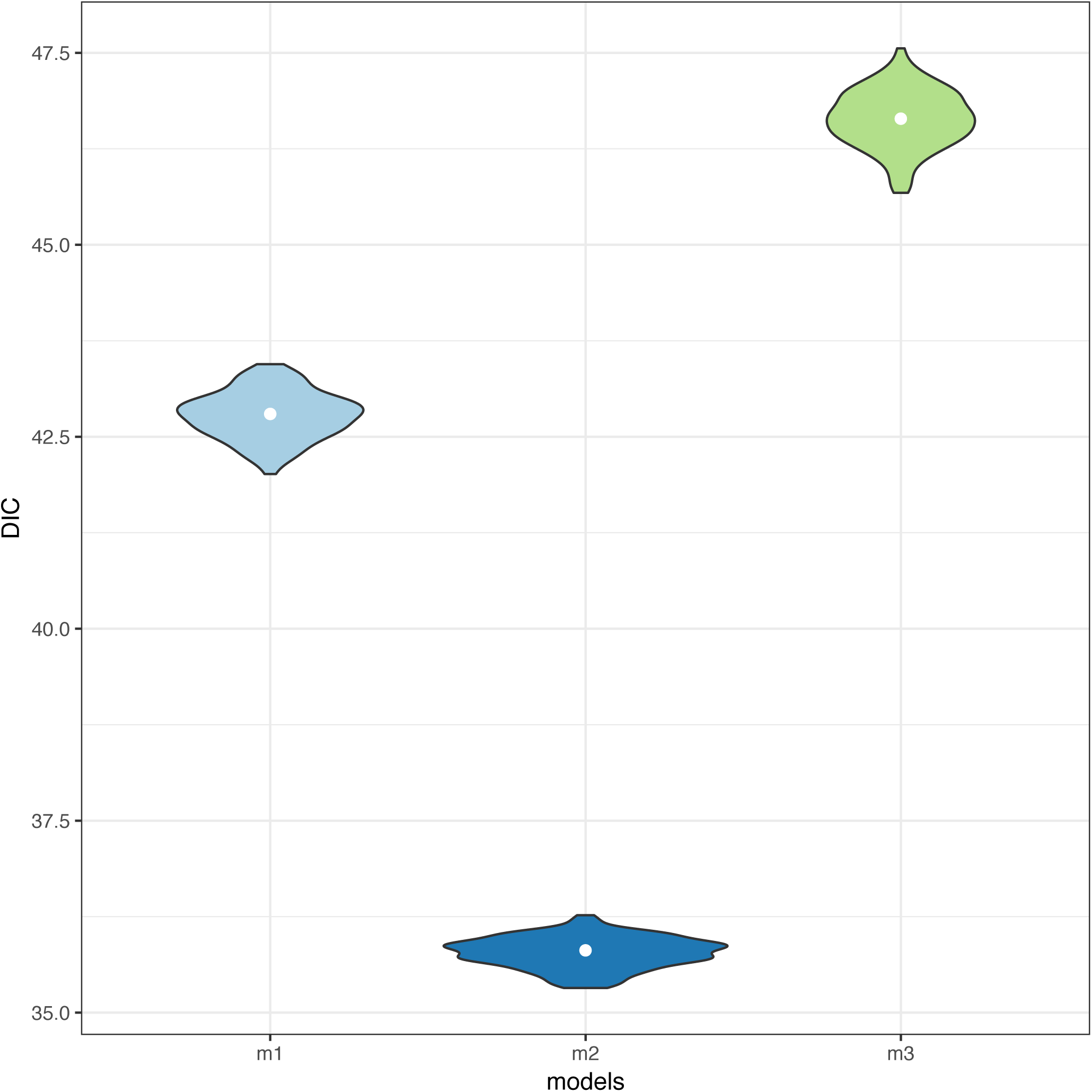
Violin-plots of the Deviance Information Criterion (DIC) of three models of the presence of S-cones as a function of plant (m1), fruit (m2), or insect (m3) diets. To account for phylogenetic variance, each model was fitted across 100 phylogenies sampled from the posterior of the phylogeny of Rojas et al. (2016).

